# Dissection of Gαs and Hedgehog signaling crosstalk reveals therapeutic opportunities to target adenosine receptor 2b in Hedgehog-dependent tumors

**DOI:** 10.1101/2025.02.21.639530

**Authors:** Sarah Krantz, Braden Bell, Katherine Lund, Natalia Salinas Parra, Yeap Ng, Natalia De Oliveira Rosa, Saikat Mukhopadhyay, Brad St Croix, Kavita Y Sarin, Roberto Weigert, Francesco Raimondi, Ramiro Iglesias-Bartolome

## Abstract

Basal cell carcinoma (BCC), the most common human cancer, is driven by hyperactivation of Hedgehog Smoothened (SMO) and GLI transcription. Gαs and protein kinase A (PKA) negatively regulate Hedgehog signaling, offering an alternative BCC development and treatment pathway. Here, using histology alongside bulk and single-cell RNA sequencing, we find that mouse BCC-like tumors that originate from Gαs pathway inactivation are strikingly similar to those driven by canonical Hedgehog SMO. Interestingly, mutations that reduce Gαs and PKA activity are present in human BCC. Tumors from Gαs pathway inactivation are independent of the canonical Hedgehog regulators SMO and GPR161, establishing them as an SMO-independent oncogenic Hedgehog signaling model. Finally, we demonstrate that activation of the Gαs-coupled adenosine 2B receptor counteracts oncogenic SMO, reducing Hedgehog signaling and tumor formation and offering a potential therapeutic strategy for BCC.

## INTRODUCTION

G-protein-coupled receptors (GPCRs) regulate somatic stem cell activity, especially in the skin, where they maintain the balance between proliferation and differentiation, and facilitate interactions between stem cells and their niche ^1^. The Frizzled class of GPCRs are central in skin stem cell biology, including Smoothened (SMO), a regulator of Hedgehog signaling, and receptors that regulate Wnt/β-catenin. In contrast to classical receptor-coupled heterotrimeric G-protein signaling, these canonical cell developmental pathways are driven by direct regulation of intracellular proteins by Frizzled class GPCRs. Hedgehog signaling is controlled by the continuous repression of SMO by patched receptors (PTCH) and is activated when secreted Hedgehog ligands bind and inhibit PTCH ^2^. Active SMO blocks the GLI-repressor SUFU, stabilizing the transcription factor GLI, which enters the nucleus and activates the transcription of target genes.

Hedgehog signaling has been involved in the development of numerous pathologies and is a primary driver of basal cell carcinoma (BCC). Mutations in PTCH and SMO are common in BCC, and overexpression of Hedgehog ligands and activated forms of SMO or GLI are sufficient to cause stem cell expansion and BCC formation in mouse skin ^3^. BCC is the most frequently diagnosed human cancer, with more than 3 million cases diagnosed each year in the US alone ^4,5^, causing a significant burden to the healthcare system. Most BCCs are easily treated by surgical removal, but a proportion of them can be challenging to treat due to their high numbers, location, or tumor spread ^6^. Alternative therapies include SMO inhibitors and immunotherapy, although side effects or therapeutic resistance undermine their use. The lack of additional targeted therapies for BCC and the fact that Hedgehog signaling is a malignancy pathway in other tumor types makes it a priority to find novel treatment options for Hedgehog-driven tumors.

Although BCC is considered to be mainly driven by alterations in canonical Hedgehog signaling, research has found a Gαs protein-dependent pathway vital to skin stem cell fate and BCC-like tumor formation ^7^. Gαs is one of the primary mediators of classic GPCR-G protein signaling, leading to the activation of adenylyl cyclases that produce the second-messenger cyclic AMP (cAMP). In mice, conditional epidermal deletion of the gene coding for Gαs (*Gnas*) is sufficient to cause an aberrant expansion of the skin basal stem cell compartment, resulting in BCC-like lesions ^7^. The effect of Gαs in tumor formation is mediated by the cAMP-regulated protein kinase A (PKA). Blocking PKA signaling by overexpression of the kinase-inhibitory domain of the PKA-inhibitor protein α (PKIα) also results in BCC-like tumors ^7^. Mechanistically, Gαs and PKA inhibition drive the activation of Hedgehog GLI and Hippo YAP1 ^7^, central mediators of tumor development in the skin. While the negative regulation of Hedgehog signaling by PKA has been established over time ^2^, this research demonstrated that Gαs or PKA inactivation is sufficient to induce BCC-like tumor formation.

The precise mechanisms that link Gαs or PKA inactivation to oncogenic Hedgehog signaling remain unclear, representing a critical gap in our understanding of BCC pathogenesis. Deciphering this relationship is essential not only to identify novel therapeutic targets for Hedgehog-driven tumors but also to expand our knowledge of GPCR-Gαs interactions in stem cell and cancer biology. Here, we present a comprehensive comparative analysis of BCC-like lesions driven by canonical oncogenic Hedgehog signaling and those caused by Gαs or PKA inactivation. Histological, bulk, and single-cell analysis reveal that BCC-like tumors from Gαs pathway inactivation are nearly indistinguishable to those arising from the canonical oncogenic Hedgehog pathway. Our results show that Gαs or PKA inactivation have the same biological outcome as SMO hyperactivity, suggesting a role for these proteins in BCC pathogenesis. Indeed, we demonstrate that mutations reducing Gαs and PKA activity are present in human BCC. A detailed analysis of GPCR networks in BCC reveals that tumors arising from Gαs pathway inactivation are independent of the canonical Hedgehog regulators SMO and GPR161. Finally, we demonstrate that activation of the Gαs-coupled GPCR adenosine 2b receptor (ADORA2B) reduces tumor formation in a BCC mouse model. Our findings emphasize both the potential and limitations of activating GPCR-Gαs signaling as a therapeutic strategy for Hedgehog-driven tumors.

## RESULTS

### G**α**s and PKA inactivation trigger oncogenic Hedgehog signaling

Gαs and PKA inactivation in mouse skin leads to highly proliferative lesions composed of basaloid cells, which form clumps and islands deeply invading the underlying stroma ^7^. These lesions were categorized as BCC-like based on several lines of evidence: lesions histologically resemble other mouse models of BCC; tumors arise primarily in the tail and ears, aligned with previous mouse models of BCC; and Gαs or PKA inactivation leads to rapid activation of Hedgehog signaling, a hallmark of BCC development. However, it remains unclear how similar these tumors are to those arising from the oncogenic activation of the canonical Hedgehog pathway. (Fig. 1A).

**Figure 1.**
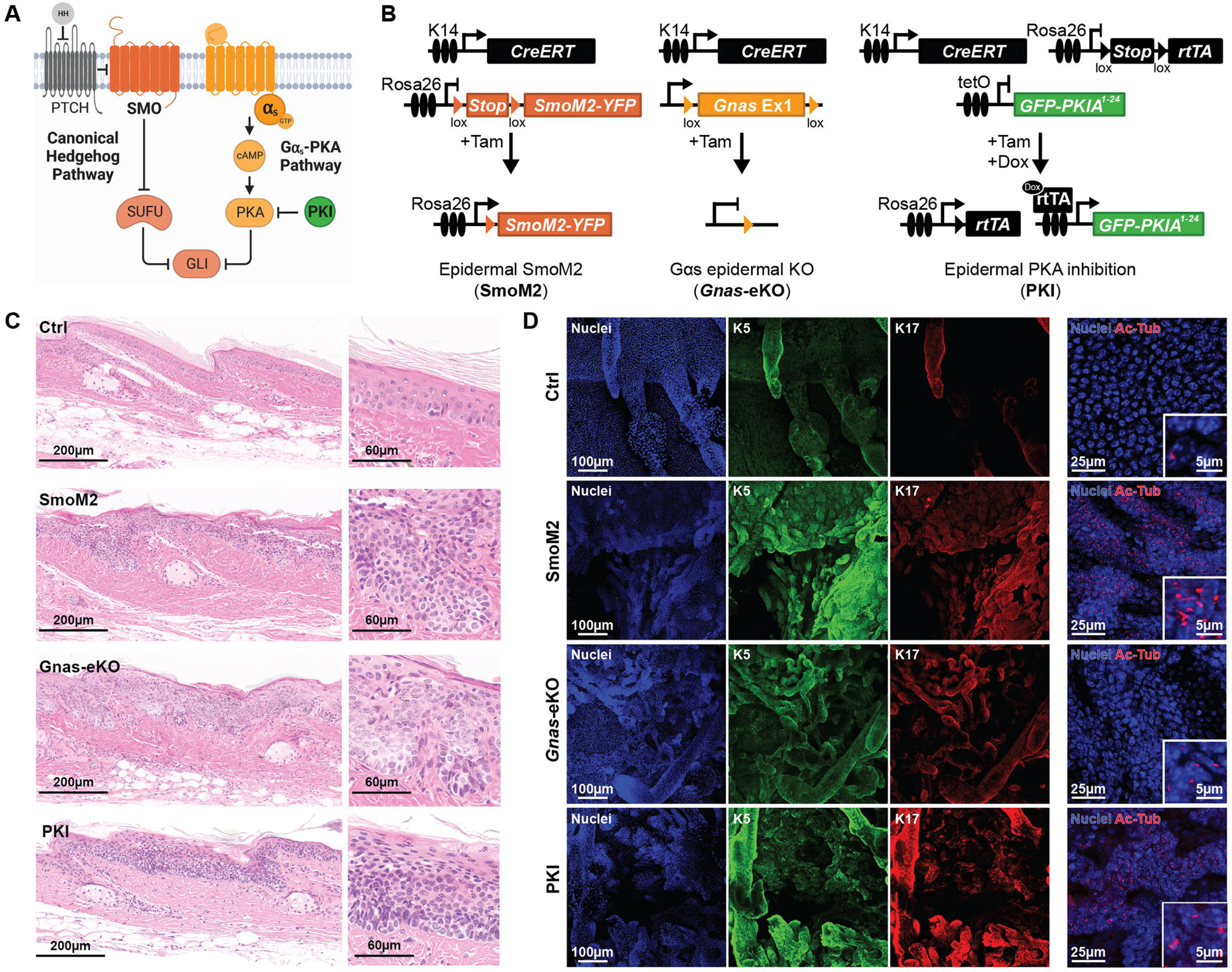
Both oncogenic Hedgehog and Gαs-PKA inactivation trigger BCC-like tumors in mouse skin. **A-** Model summarizing the canonical Hedgehog pathway and the Gαs-PKA pathway, which regulate GLI transcription. **B-** Schematic representations of the SmoM2, *Gnas*-eKO, and PKI inducible mouse models. **C-** Representative images from H&E staining of tail skin from control wild type (Ctrl), SmoM2, Gnas-eKO, and PKI mice showing similar BCC tumors with magnification of the epidermis. **D-** Representative images from immunofluorescence (IF) staining of tail skin whole mounts showing expression of basal marker keratin 5 (K5), the BCC marker keratin 17 (K17), and the cilia marker acetylated tubulin (Ac-Tub).

To compare tumors arising from canonical Hedgehog signaling versus Gαs pathway inactivation, we utilized a series of animal models in which inducible Cre recombination is driven by the keratin 14 promoter (K14CreERT), targeting the epidermal basal proliferative compartment of the skin ^8^ (Fig. 1B). To trigger canonical oncogenic Hedgehog signaling we took advantage of a mouse model that conditionally expresses constitutively active SMO (SmoM2) under the ROSA26 promoter upon Cre recombination ^9^ (abbreviated SmoM2). The SmoM2 mouse model represents SMO-inhibitor-resistant BCC ^10^. Gαs pathway inactivation was carried out either by knockout of the *Gnas* gene, utilizing mice carrying loxP sites surrounding *Gnas* exon one ^11^ (abbreviated *Gnas*-eKO), or by overexpression of the kinase-inhibitory domain of PKIα, with mice carrying LoxStopLox-rtTA ^12^ and tetracycline-inducible PKIα1-24 peptide tagged with GFP, as previously described ^7^ (abbreviated PKI). Between 3 to 6 weeks after tumor induction, all three models developed thickening of the epidermis, primarily on ears, snout, and paws. These thickened skin areas presented basaloid tumor lesions that invaded the underlying dermis, morphologically resembling superficial and nodular BCC ^7,9,13^ (Fig. 1C). Immunofluorescence staining of tail epidermis whole mounts revealed that the BCC-like lesions in the three mouse models were positive for the basal keratinocyte marker keratin 5 (K5) and the basaloid skin tumor marker keratin 17 ^14^ (K17, Fig. 1D). Tumor cells in all three models were ciliated, as evidenced by staining with the cilia marker acetylated tubulin (Ac-Tub, Fig 1D. Cilia are uncommon and only present close to hair follicles in control mice).

Bulk RNA-sequencing (RNAseq) of tail keratinocytes of BCC mice revealed similar changes in gene expression among the mouse models, particularly between SmoM2 and *Gnas*-eKO (Fig. 2A and S1A). Common genes significantly dysregulated showed enrichment for pathways related to BCC, including Hippo, TGF-beta, and Hedgehog signaling (Fig. 2B). Indeed, numerous genes associated with Hedgehog signaling were upregulated in the skin, including *Gli* and *Ptch* (Fig. S1B). Functional analysis of transcriptional regulators through Ingenuity Pathway Analysis (IPA) indicated upregulation of BCC-related transcriptional networks (Fig. S1C).

**Figure 2.**
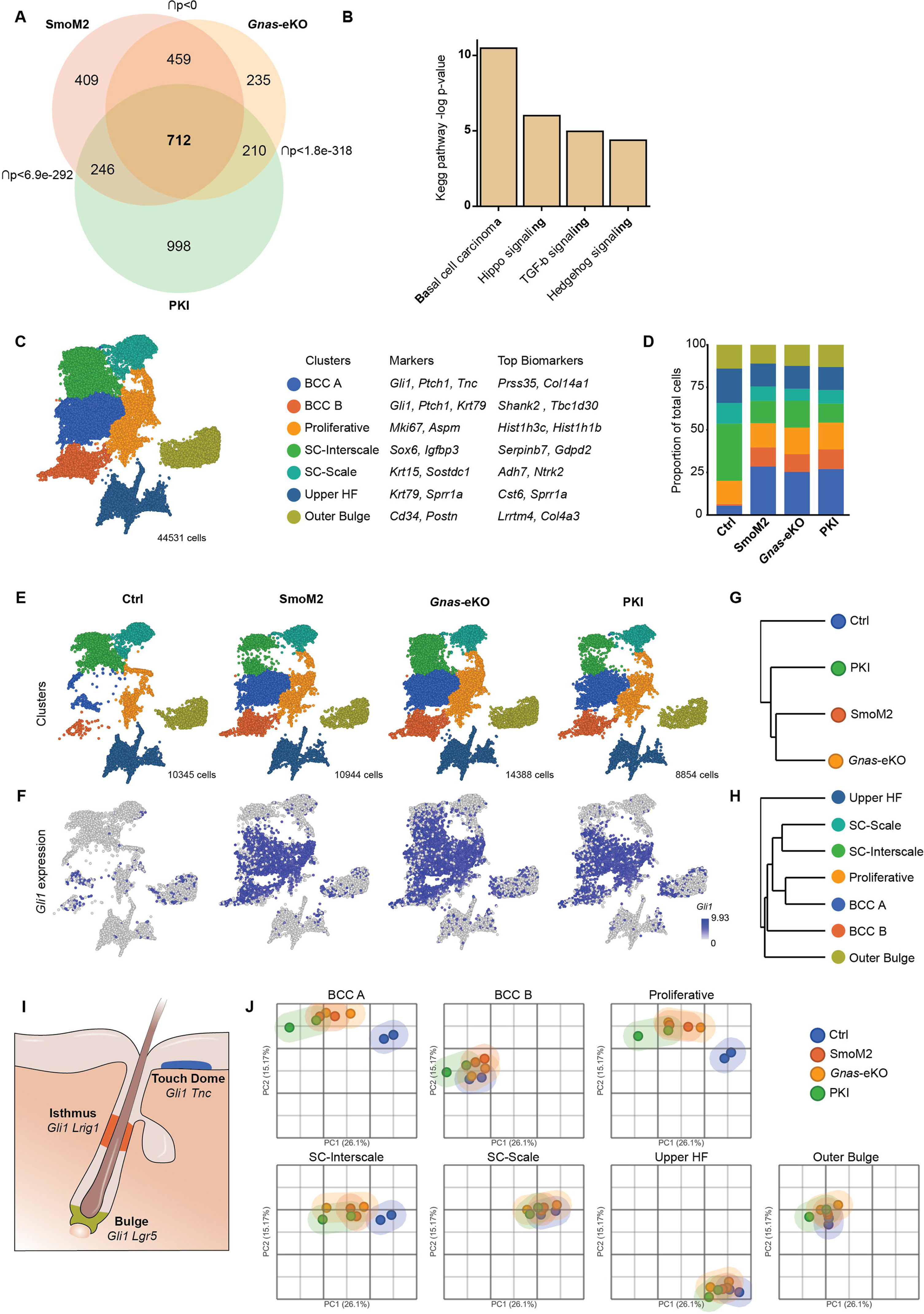
SMO and Gαs/PKA inactivation trigger oncogenic Hedgehog signaling. **A-** Venn diagram showing the differentially regulated genes overlap between mouse models by bulk mRNA-sequencing (q < 0.05, |FC| ≥ 1.5). p indicates the p-value of the overlap, Fisher’s exact test. N= 4 of each SmoM2, Gnas-eKO, and PKI mice; and N=9 control mice. **B-** Graph indicating the enriched Kegg pathway terms in the 712 overlapping genes shown in (A). **C-** Uniform Manifold approximation and projection plot (UMAP) of cell clusters from all mouse models from tail single-cell sequencing. Gene markers used for cell identification and top enriched biomarkers for each cluster are indicated. SC=stem cell, HF= hair follicle. N= 2 mice per genotype, 44531 total cells. **D-** Relative proportion of each cell population shown in C for each mouse model. **E-** UMAP of cell clusters for each mouse models from tail single-cell sequencing. N= 2 mice per genotype. Number of cells sequenced for each genotype is indicated. **F-** Expression of *Gli1* projected into each UMAP. **G and H-** Unsupervised hierarchical clustering of pseudobulk single-cell RNAseq data based on mouse genotype (G) or cell cluster (H). **I-** Schematic showing Gli+ stem cell compartments in skin. **J-** Principal component analysis plots (PCA) showing clustering of the different cell populations for each mouse. Calculated with pseudobulk single-cell RNAseq data based on mouse genotype and cell cluster. See also Figure S1.

We performed single-cell RNA-sequencing of tail keratinocytes from our animal models to better understand the changes triggered by oncogenic Hedgehog signaling. Tail skin cell populations were classified by unsupervised clustering and expression of known markers for each cell type ^15–18^ (Fig. 2C, S1D and E). Only basal cells were selected for further analysis. Interfollicular stem cells from the mouse tail were divided into two different populations, depending on scale or interscale markers ^17^, named SC-Interscale (*Sox6* and *Igfbp3*+) and SC-Scale (*Krt15* and *Sostdc1*+). BCC-like cells were identified based on the presence of Hedgehog signaling markers (*Gli1-2-3*, *Ptch1-2, Smo,* and *Gas1*) and increased cell number for that population in our BCC models with respect to control mice (Fig. 2D). We were able to confirm that cells in *Gnas*-eKO mice have reduced *Gnas* expression and that *Gnas* depletion is enriched in the BCC cell populations (Fig. S1F).

Remarkably, SmoM2, *Gnas*-eKO, and PKI mice presented highly similar cell profiles and upregulation of Hedgehog targets (Fig. 2E, F, and G). We found that BCC-like cells clustered in two separate populations, BCCA and BCCB, that matched to cell clusters present in control mice (Fig. 2E). The BCC-like cells in control mice corresponded to *Gli*+ populations (Fig. 2E and F). The gene expression profile of BCCA cells was more closely related to the proliferative compartment and basal stem cells (Fig. 2H), possibly indicating a higher proliferative potential for these cells. GLI is present in three main stem cell areas in normal skin (Fig. 2I). Based on markers for these GLI+ stem cells, we hypothesize that BCCA cells correspond to a touch-dome stem cell-like state (*Tnc*+), while BCCB matches an isthmus stem cell phenotype (*Krt79*+), although both populations present broad expression of hair follicle stem cell markers like *Lgr5* and *Lrig1* (Fig. S1D and E).

The populations with more changes in their RNA profile corresponded to BCC-like cells, proliferative compartment, and interscale stem cells (SC-Interscale) based on clustering analysis of pseudobulk samples (Fig. 2J). This was not necessarily a reflection of changes in the proportion of total cells for those corresponding populations since the BCC-like cells expanded in SmoM2, *Gnas*-eKO, and PKI mice, with a decrease in the number of SC-Interscale, while the proportion of Proliferative cells did not change (Fig. 2D).

Overall, our results establish that tumor cells arising from Gαs pathway inactivation are almost indistinguishable from those resulting from canonical oncogenic Hedgehog pathway activation. We also find that BCC-like tumor cells have gene expression profiles similar to those of touch dome and isthmus basal stem cells. Remarkably, the main gene expression changes triggered by oncogenic Hedgehog signaling are reflected in the BCCA and proliferative cells, while the oncogenic activation does not significantly alter other stem cell compartments.

### Hair follicle stem cells have limited potential to generate BCC-like tumors

Bulk and single-cell RNAseq confirmed the similarity of BCC-like lesions to hair follicle stem cells, suggesting a hair follicle origin for these tumors. However, evidence suggests that by reprogramming towards a hair follicle phenotype ^19^, BCC could originate from other keratinocyte stem cell populations. As indicated above, BCC cells expressed a mix of stem cell markers, although they principally match touch-dome and isthmus stem cell-like states. To better characterize the origin of BCC-like cells, we analyzed the tumorigenic potential of different hair follicle stem cell populations by crossing our inducible PKI transgene with drivers specific for isthmus *Lrig1* ^20^ or bulge *Lgr5* ^21^ hair follicle stem cells and tracked tumor formation over time (Fig. 3A and B). Remarkably, *Lgr5* cells were not able to drive tumor formation (Fig. 3C). In contrast, *Lrig1* cells formed lesions limited to the mid-hair follicle area (Fig. 3C). These *Lrig1*-derived tumors did not migrate into the interfollicular epidermis or invade the dermis, as seen with the K14 driver. We validated by immunofluorescence that PKI is expressed in the specific hair follicle compartments by staining tail epidermis whole mounts against GFP (Fig. 3D), which is fused to PKI in our transgenic mouse model. Our results indicated that *Lrig1* and *Lgr5* hair follicle stem cells are unlikely to be the primary origin for BCC-like tumors following PKA inactivation.

**Figure 3.**
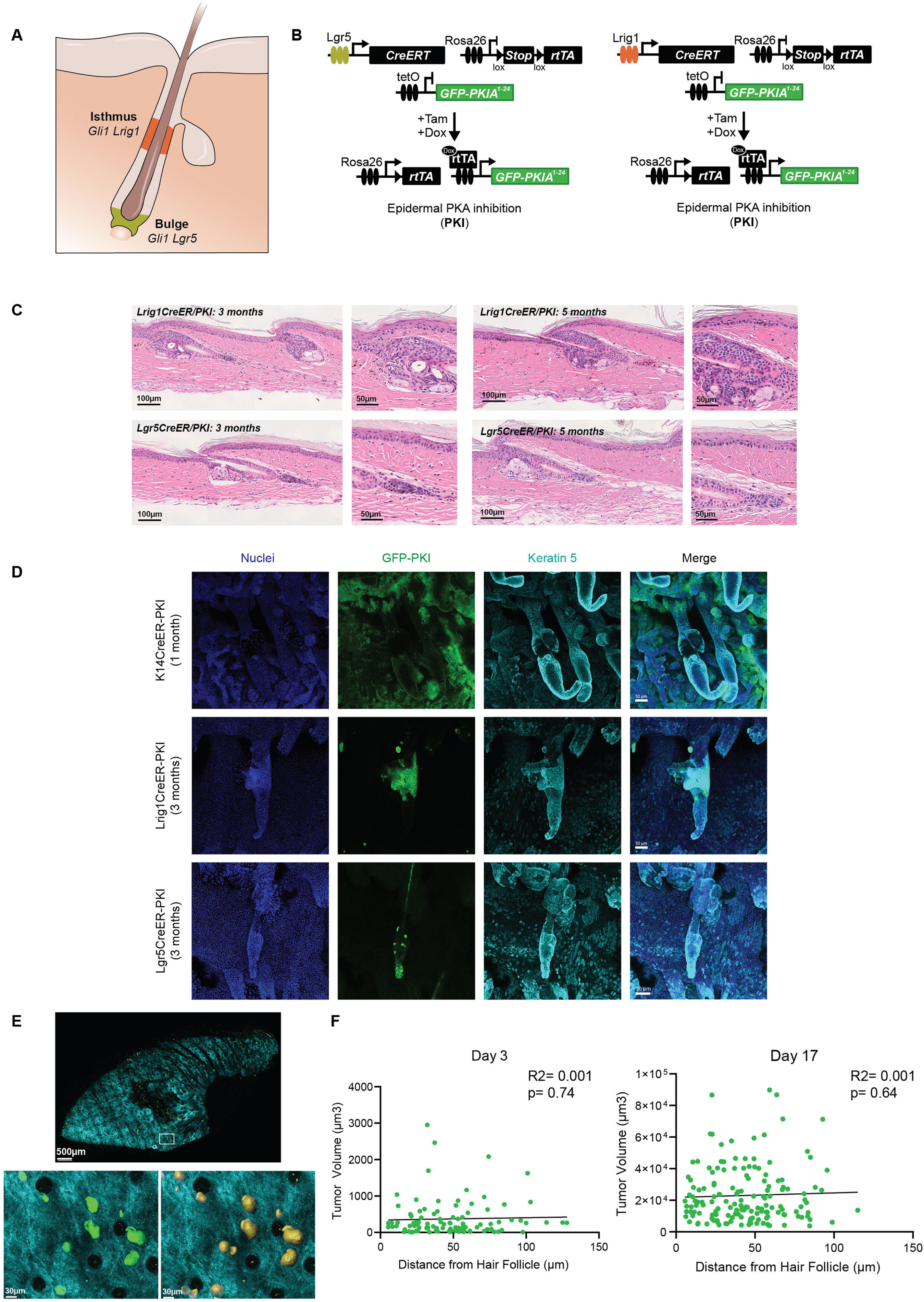
Hair follicle stem cells have limited potential to generate BCC-like tumors. **A-** Schematic showing bulge and isthmus stem cell compartments in skin. **B-** Schematic representations of the mice used to target PKA inactivation to hair follicle stem cell compartments. **C-** Representative images from H&E staining of tail skin from the indicated mice three and five months after induction. **D-** Representative images from IF staining of tail skin whole mounts showing expression of PKI and basal marker K5, at the indicated time after induction. **E-** Two-photon image of mouse whole ear with GFP-PKI tumors (green) and collagen (cyan) visualization from a mouse 17 days after induction. Magnified images show GFP-PKI tumors. Hair follicles can be visualized by disruptions in the collagen (black circular areas). On the right, a rendering for tumor volume analysis (IMARIS) used for quantification is shown. **F-** Quantification of the distance between tumors and closest hair follicle. Each dot represents an individual tumor. Best fit regression line is shown. Day 3, N= 98 tumors from 3 mice; day 17, N=155 tumors from 3 mice.

To better understand tumor origin and evolution in our BCC model, we took advantage of the GFP fluorescent protein fused to PKI to track BCC-like tumors by two-photon microscopy in K14-driven mice. Utilizing second-harmonic generation to image collagen ^22^, we can also observe the tissue structure and track the location and growth of tumors according to their distance from the hair follicle (Fig. 3E and F). Our results indicate that tumor formation (day 3) and tumor growth (week 2) have no significant correlation from the distance to the hair follicle (Fig. 3F), supporting that despite their hair follicle stem cell phenotype, BCC-like cells may not originate from hair follicles themselves.

### G**α**s and PKA inactivating mutations are present in human BCC

The similarity between our tumor models suggested that genomic alterations in Gαs-PKA signaling members could contribute to the genetic risks of Hedgehog-driven tumors. Indeed, alterations in cAMP-dependent pathways, *Gnas*, and PKA have been found in Hedgehog-driven medulloblastoma ^23,24^, although this has not been explored in detail for human BCC. Analysis of a tumor dataset of human BCC tumors ^25^ shows mutations in genes coding for Gαs and PKA (Fig. 4A).

**Figure 4.**
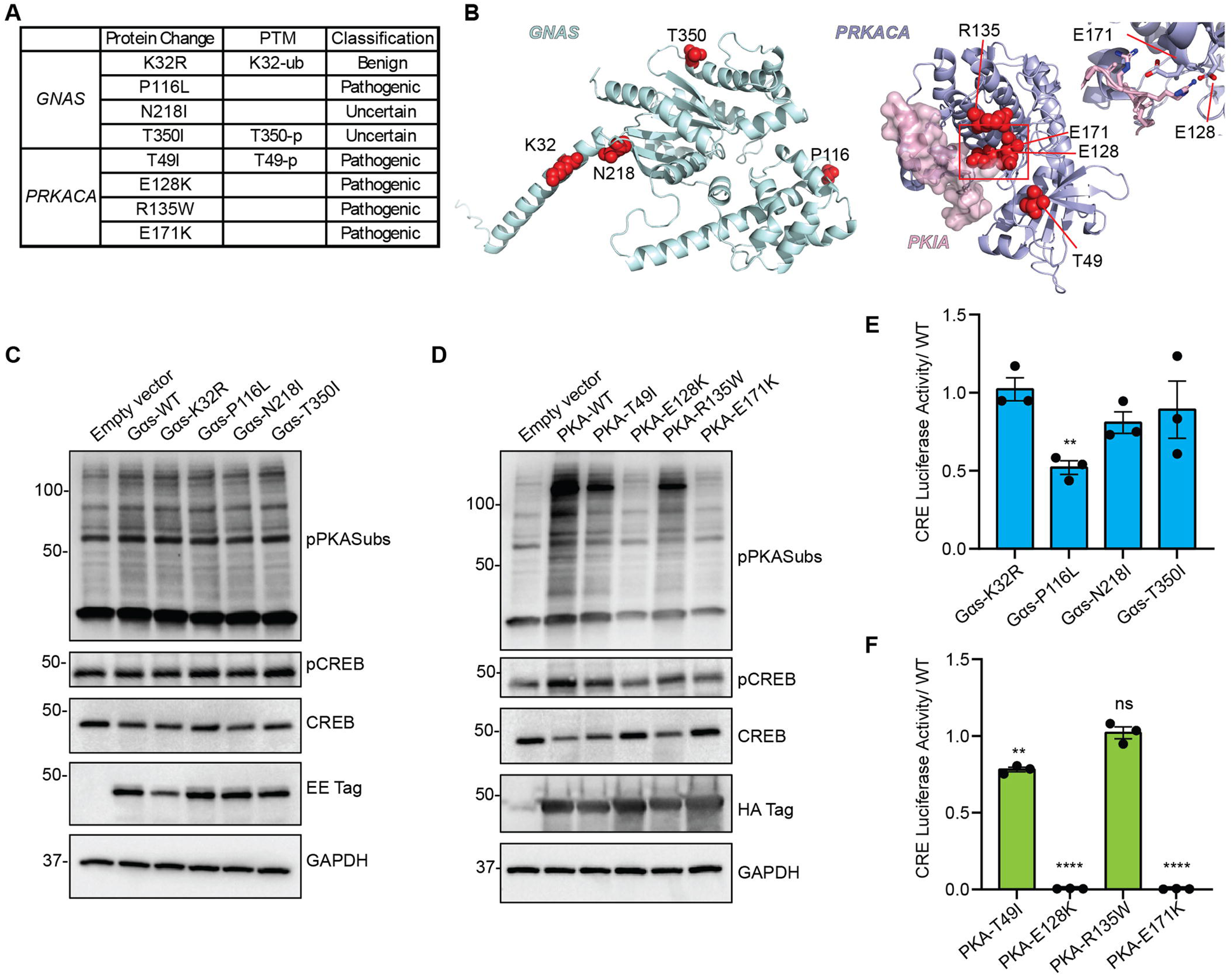
Gαs and PKA inactivating mutations are present in human BCC. **A-** Table of Gαs (*GNAS*) and PKA catalytic alpha subunit (*PRKACA*) mutations detected in human BCC, indicating post-translational modifications (PTM) at those sites and classification of predicted mutation effect. **B-** Protein models of GNAS and PRKACA with mutated amino acids highlighted in red. PRKACA is shown in complex with the pseudosubstate PKIA. Magnification on the right shows amino acids important for substrate interaction between PKA and PKI. **C and D-** Western blot analysis of expression of EE-tagged Gαs (C), HA-tagged PKA (D), wild type (WT), or mutants, and downstream signaling pathway components in transfected HEK293 cells. pPKA-subs= Phospho-PKA Substrate RRXS*/T*. Molecular weight markers (kDa) are indicated on the left. **E and F-** Transcriptional activity of CREB measured by CRE-luciferase assay, in HEK293 cells transfected with Gαs (E) or PKA (F) variants. Graphs show mean ± SEM. In E, N=3, one sample Wilcoxon test; in F, N=3, one sample Wilcoxon test.

If Gαs and PKA inactivation participate in BCC development, it would be expected that some of these mutations will decrease Gαs and PKA activity. Mapping and effect predictor analysis of Gαs and PKA mutations indicated potential altered protein activity for some variants (Fig. 4A and B). Notably, two mutations in the PKA catalytic subunit alpha (*PRKACA*) are in residues that are essential for the kinase to interact with substrates (E128 and E171, Fig. 4B). To test the biological effect of the mutations, we measured the activity of Gαs and PKA variants present in BCC. All mutants were expressed at similar levels as wild-type (WT) counterparts (Fig. 4C and D). While Gαs mutants showed similar activity levels to WT, all PKA mutations reduced phosphorylated PKA substrates and CREB phosphorylation (Fig. 4C and D). To expand these results, we next utilized a CRE-luciferase reporter that indicates the activity of the cAMP-dependent pathway by measuring CREB transcription. Interestingly, one *GNAS* variant, Gαs-P116L showed reduced CREB reporter activation (Fig. 4E). The *PRKACA* variants PKA-E128K and E171K showed complete abolishment of activation of CREB reporter and PKA-T49I showed reduced activity (Fig. 4F). These results show that disruptive mutations in Gαs and PKA are present in human BCC samples, indicating that inactivation of this pathway probably contributes to human BCC.

### BCC tumors arising from G**α**s pathway inactivation are independent of the canonical Hedgehog regulators SMO and GPR161

Our results indicate that Gαs and PKA are necessary to block Hedgehog signaling during normal epithelial differentiation, suggesting that potential Gαs-coupled GPCRs are involved in the regulation of stem cell fate in the skin. Analysis of gene expression data from our bulk and single-cell data identified numerous GPCRs expressed in tumor cells from our mouse models (Fig. 5A and S2). Interestingly, GPCRs that regulate Hedgehog signaling, like *Smo* and *Gpr161*, are upregulated in BCC-like cells (Fig. S3A).

**Figure 5.**
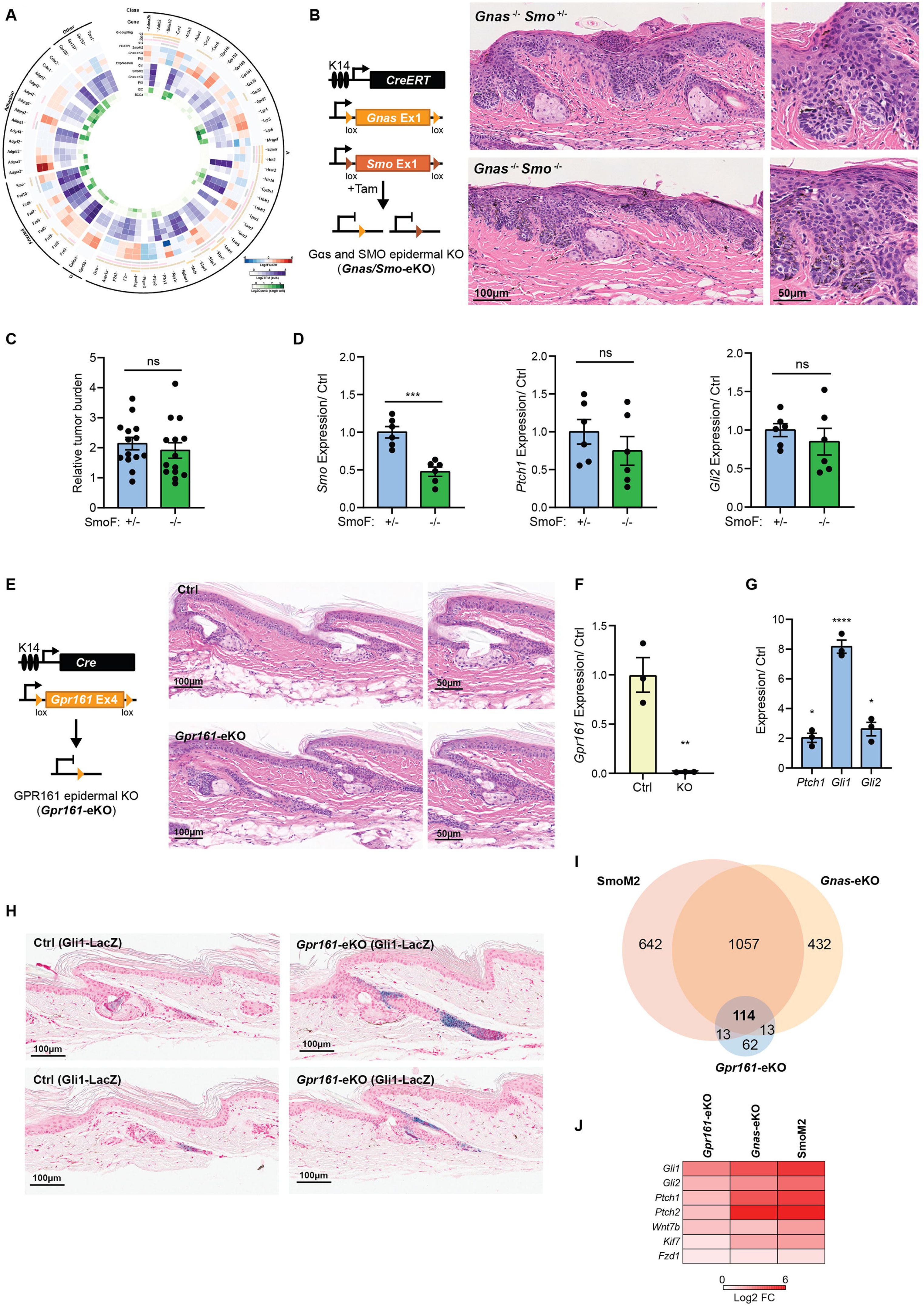
BCC tumors arising from Gαs pathway inactivation are independent of canonical Hedgehog regulators. **A-** Circos plot representing BCC GPCR expression in mice, showing class, gene name, Gα coupling, fold change (FC) and expression level (Log2TPM) in bulk RNAseq, level of expression in single-cell RNA sequencing (Log2Counts) in interscale (iSC) and BCCA clusters. See the amplified image in Figure S2. **B-** Schematic representation of the mouse model used to target Gαs and SMO deletion to the skin. Images on the right show representative H&E staining of tail skin from the indicated mice 33 days after induction. -/+ indicates heterozygous knockout and -/- indicates homozygous knockout. **C-** Quantification of tumor burden per mouse in H&E tail skin stainings in *Gnas* knockout mice with *Smo* heterozygous (+/-) or homozygous (-/-) knockout. Graphs show mean ± SEM. N= 14 SmoF +/- mice, N= 14 SmoF -/- mice, t-test. **D-** qRT-PCR analysis of mRNA expression of indicated markers in tail epidermal keratinocytes isolated from *Gnas* knockout mice with *Smo* heterozygous (+/-) or homozygous (-/-) knockout 33 days after induction. Graphs show mean ± SEM. N= 6 SmoF +/- mice, N= 6 SmoF -/- mice, t-test. SmoF +/- mice were used as control (Ctrl). **E-** Schematic representation of the mouse model used to target GPR161 deletion to the skin. Images on the right show representative H&E staining of tail skin from the indicated mice five weeks after induction. **F and G-** qRT-PCR analysis of mRNA expression of indicated markers in tail epidermal keratinocytes isolated from 4-month-old control (Ctrl) or *Gpr161*-eKO mice. Graphs show mean ± SEM. In F and G, N=3 control mice and N=3 *Gpr161*-eKO mice; F, t-test; and G, one sample Wilcoxon test. **H-** Representative image of βGal staining in tail skin from 12-week-old control (Ctrl) and *Gpr161*-eKO (K14-Cre) mice showing GLI^+^ cells. **I-** Venn diagram showing the differentially regulated genes overlap between indicated mouse models by bulk mRNA-sequencing (q < 0.05, |FC| ≥ 1.5). SmoM2 and *Gnas*-eKO are K14-CreERTM as shown in Figure 2A; and *Gpr161*-eKO are from K14Cre 4-month-old mice, N= 3 control and 3 knockout mice. **J-** Heat map from bulk mRNA-sequencing data comparing the fold change (log2FC) increased in Hedgehog signaling targets in the indicated mice. See also Figures S2, S3 and S4.

Although it is known that Gαs and PKA act downstream of SMO to regulate the Hedgehog pathway ^2^, one open question is whether oncogenic Hedgehog signaling following Gαs and PKA inactivation depends on SMO. To test this possibility, we performed a skin-specific double knockout of *Smo* and *Gnas*, or knockout of *Smo* in the context of PKI expression (Fig. 5B and S3B). Surprisingly, the knockout of *Smo* did not alter tumor formation or Hedgehog signaling activation in *Gnas*-eKO or PKI mice (Fig. 5B and C; Fig. S3B and C). Gene expression analysis of tail keratinocytes from these mice indicated a significant reduction in the expression of *Smo* but not in Hedgehog target genes like *Gli2* or *Ptch1* following *Smo* knockout (Fig. 5D and S3D).

Combined with our previous study demonstrating that Gαi-GPCR activation does not lead to BCC ^26^, our *Smo* knockout results suggested that a Gαs-coupled receptor must be present in the skin, which inactivates Hedgehog signaling independently from SMO during normal homeostasis. We hypothesized that if the effect is mediated by a single GPCR, skin-specific knockout of this Gαs-coupled receptor would phenocopy *Gnas*-eKO BCC formation. A literature review of skin-specific or complete knockouts of the GPCRs expressed in BCC cells did not identify any receptor whose absence results in BCC-like lesions. However, for two of these GPCRs, *Gpr161* and *Adgra2* (also known as *Gpr124*), their complete knockout is embryonically lethal ^27,28^, and the specific skin knockout has not been analyzed. *Adgra2* is expressed at very low levels in the skin of control mice while is highly upregulated in our BCC mouse models (Fig. S3A). *Adgra2* skin-specific knockout mediated by constitutive CRE recombination driven by the keratin 14 promoter (K14Cre ^8^) and lox-p sites on *Adgra2* exon 1 ^28^, did not affect skin homeostasis (Fig. S4A, B and C).

We next moved into analyzing GPR161, a Gαs-coupled receptor that is one of the primary inhibitors of Hedgehog signaling during development and tumor formation ^2,27,29^. To confirm that GPR161 has similar functions in keratinocytes, we generated inducible-keratinocyte cell lines overexpressing wild type (WT) or V129E GPR161, a mutation that abolishes GPR161 interaction with Gαs ^27^. Overexpression of WT but not mutant GPR161 leads to the expression of GLI3-repressor and activation of CREB (Fig. S4D and E), indicating that GPR161 can regulate Gαs and Hedgehog signaling in keratinocytes. Surprisingly, however, inducible (K14CreERT) or constitutive (K14Cre) skin-specific knockouts utilizing a floxed-*Gpr161* mouse ^30^ did not show any phenotypic or histological alterations (Fig. 5E and F; Fig. S4F, G and H).

Analysis of the expression of Hedgehog targets in *Gpr161* knockout tail skin keratinocytes indicated that, despite the lack of phenotype, keratinocytes show increased expression of Hedgehog targets (Fig. 5G). Utilizing *Gli1* β-galactosidase reporter mice (*Gli1^lacZ^*) ^31^, we were able to visualize that *Gpr161* epithelial knockout led to an increase in *Gli1*+ cells limited to the hair follicle (Fig. 5H), without leading to ectopic activation of *Gli1* as it has been shown for *Gnas*-eKO mice ^7^. Comparison of bulk RNAseq analysis of *Gpr161* tail skin keratinocytes with SmoM2 and *Gnas*-eKO mice indicated that fewer genes are differentially regulated by *Gpr161* knockout, although several of these genes were shared across the mouse models (Fig. 5I). Comparing the expression of Hedgehog targets showed that *Gpr161* epithelial knockout leads to a less pronounced increase in these genes, compared with SmoM2 and *Gnas*-eKO (Fig. 5J). Our results show that *Gpr161* knockout in basal keratinocytes leads to a non-oncogenic increase in Hedgehog signaling, suggesting that GPR161 only partially controls the Hedgehog pathway in the skin.

Overall, our GPCR analysis indicates that the tumor-suppressive function of Gαs is probably not linked to a single Gαs-coupled GPCR but might involve multiple upstream receptors. On the other hand, our results demonstrate that BCC tumors arising from Gαs pathway inactivation are independent of the canonical Hedgehog regulators SMO and GPR161 and establish *Gnas*-eKO/PKI models as a unique resource to understand BCC arising from SMO-independent Hedgehog signaling.

### Activation of G**α**s-coupled adenosine receptor 2b reduces BCC tumor formation

Since Gαs and PKA are essential inhibitors for the Hedgehog pathway in the skin, elevating cAMP levels and activating PKA could be a viable option to block oncogenic Hedgehog signaling (Fig. 6A). Inhibitors of phosphodiesterases (PDE), the enzymes mediating degradation of cAMP, can suppress Hedgehog signaling and medulloblastoma growth in mouse models ^32^. However, we found that systemic delivery of the PDE4 inhibitor rolipram, the PDE7 inhibitor BRL50481, or their combination, did not affect the growth of SmoM2-induced tumors (Fig. 6B and C).

**Figure 6.**
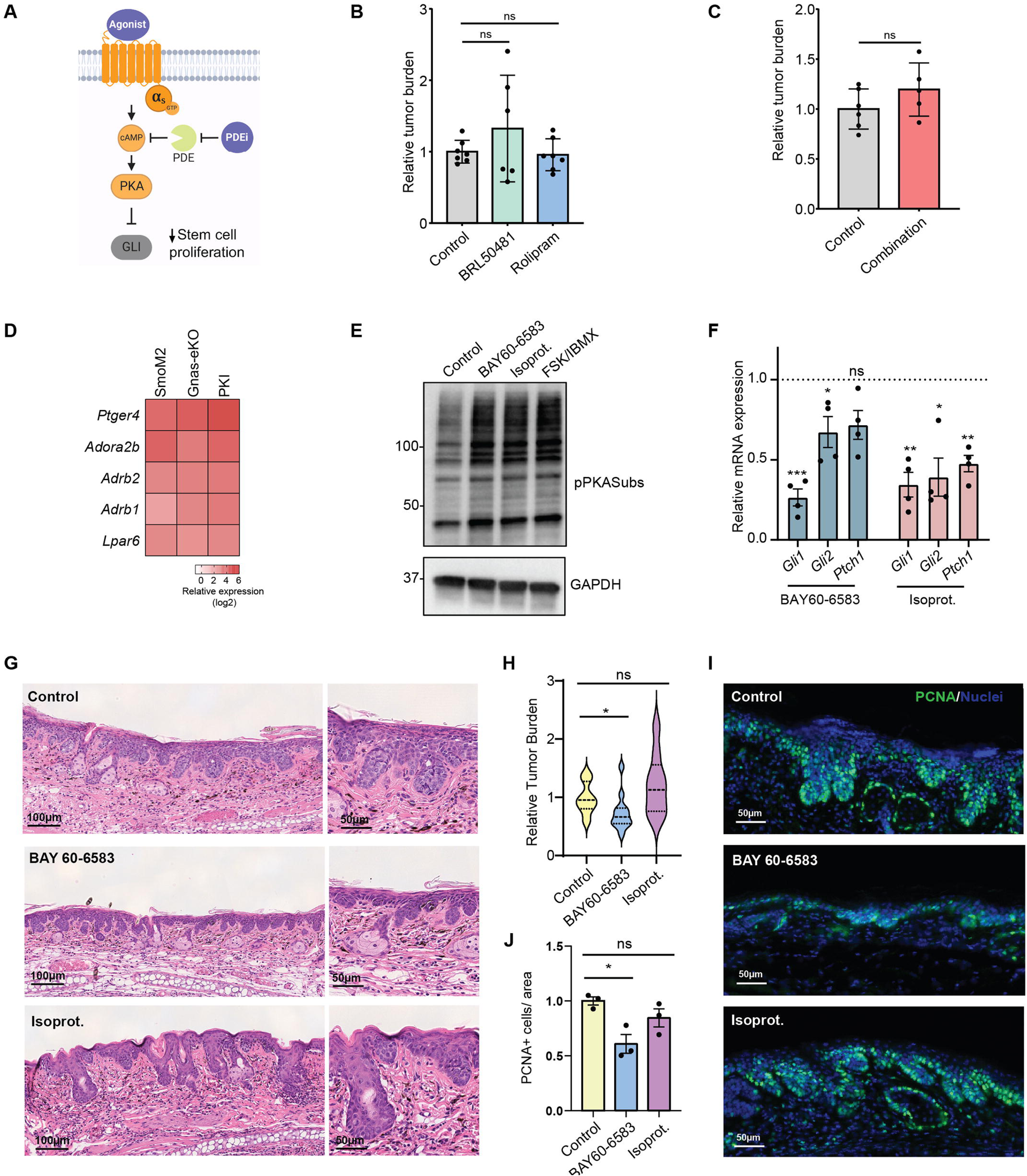
Activation of Gαs-coupled receptors reduces BCC tumor formation. **A-** Model summarizing the Gαs-PKA pathway and potential therapeutic interventions to reduce oncogenic Hedgehog signaling, including using phosphodiesterase inhibitors (PDEi) to block cAMP degradation or Gαs-coupled GPCR agonists to increase pathway activation. **B and C-** Quantification of tumor burden per mouse in H&E tail skin from SmoM2 mice treated with the PDEi BRL50481 or rolipram (B) or a combination of both (C). Graphs show mean ± SEM. For B, N= 7 control, 6 BRL50481, 7 rolipram. treated mice, Anova with t-test; For C, N= 6 control, 5 combination treated mice, Anova with t-test. **D-** Heat map from single cell RNAseq data from BCCA cells comparing the relative expression of the indicated Gαs-coupled GPCRs. **E-** Western blot analysis of phospho-PKA substrates (RRXS*/T*, pPKASubs) in a cell line derived from SmoM2 BCC tumors showing activation of PKA by treatment with BAY60-6583 or isoproterenol (Isoprot.) for 15 minutes. Forskolin and IBMX (FSK/IBMX) treatment is used as a positive control. Molecular weight markers (kDa) are indicated on the left. **F-** qRT-PCR analysis of mRNA expression of indicated markers in a SmoM2 cell line treated with the indicated drugs for 24hs. Graphs show mean ± SEM. N= 4 samples per condition, one sample Wilcoxon test. Dotted line indicates expression level of control samples. **G-** Representative images from H&E staining of ear skin from SmoM2 mice treated topically for 10 days with the indicated drugs or vehicle (Control). **H-** Violin plot depicting distribution of tumor burden of mice shown in (G). N= 15 control, 12 BAY60-6583, 9 isoprot. treated mice, Welch’s t-test. **I and J-** IF (I) and quantification (J) of the proliferation marker PCNA in mice shown in (G). N= 3 mice for each condition, one sample Wilcoxon test. See also Figure S5.

Another potential way to increase cAMP in cells is to activate Gαs-coupled GPCRs (Fig. 6A). Analysis of gene expression in our single-cell BCC data revealed several targetable Gαs-coupled GPCRs present in tumor cells, including prostaglandin E receptor 4 (*Ptger4*), adenosine receptor 2b (*Adora2b*) and β-Adrenergic Receptors (*Adrb1-2*), among others (Fig. 6D). As a proof of concept to induce Gαs-coupled GPCR signaling and reduce Hedgehog activity, we utilized BAY60-6583 ^33^, a specific agonist for ADORA2B, and isoproterenol, an agonist for β-adrenergic receptors. Treatment of a cell line generated from SmoM2 mice that has BCC-like characteristics (Fig. S5A to F) showed activation of PKA signaling by treatment with BAY60-6583 or isoproterenol (Fig. 6E). Furthermore, this treatment led to reduced expression of Hedgehog signaling targets like *Gli1* and *Gli2* (Fig. 6F).

Our results indicated that activation of PKA by Gαs-coupled GPCRs could be used to reduce Hedgehog activation. To demonstrate this in an in vivo setting, we treated SmoM2 mice with daily skin topical application of BAY60-6583 or isoproterenol for 10 days, two weeks after tumor induction. Surprisingly, BAY60-6583 reduced tumor burden, while isoproterenol did not significantly alter tumor growth (Fig. 6G and H). BAY60-6583 led to a significant reduction in tumor cell proliferation (Fig. 6I and J). Although the base for the discrepancy between BAY60-6583 and isoproterenol effects is unclear, one potential explanation is the differential expression of their respective receptors across stem cell compartments in the epidermis (Fig. S5G). In addition, *Adora2b* expression is increased in tumor cells in BCC mice when compared to control mice, while β-adrenergic receptor expression is reduced (Fig. S5G). Both our in vitro and in vivo data support that activating Gαs-coupled GPCRs can inhibit oncogenic Hedgehog signaling.

## DISCUSSION

Oncogenic and dysregulated Hedgehog signaling contributes to numerous human pathologies, including cancer. Here, we dissect the role of Gαs and PKA in the regulation of mammalian Hedgehog, and we demonstrate that inactivation of Gαs and PKA is sufficient to trigger oncogenic Hedgehog signaling at the same level and with the same biological consequences as with constitutive active SMO. We also determine that tumors arising from Gαs pathway inactivation are independent of the canonical Hedgehog regulators SMO and GPR161 and establish the *Gnas*-eKO and PKI models as a unique resource to understand tumor formation arising from noncanonical activation of Hedgehog signaling. Our findings expand the roles of Gαs and PKA in Hedgehog signaling and highlight the crucial function of Gαs and Gαs-coupled receptors in regulating this essential developmental pathway in stem cell biology. The GPCR-Gαs axis is the target of many commonly used drugs in humans, including adrenergic drugs and agonists for glucagon-like peptide receptors, so identifying how Gαs and its signaling partners regulate stem cell pathways is crucial to understand the impact of these treatments in tissue homeostasis and cancer.

Our results indicate that continuous activation of Gαs and PKA is essential to maintain normal levels of Hedgehog activity in the skin stem cell compartment and that this function is epistatic to SMO, supporting the idea that SMO primarily regulates the inhibitory effect of PKA on Hedgehog signaling. Indeed, SMO activity is believed to be predominantly mediated by direct inhibition of PKA through a pseudosubstrate site ^34^ and Hedgehog pathway hyperactivity is independent of SMO in other models of cAMP signaling inactivation ^27,35,36^, highlighting the overlapping relationship between Gαs, PKA, and Hedgehog signaling.

BCC is a prototypical GPCR-driven cancer and an excellent model to study how imbalanced signaling leads to dysregulation of normal cycles of stem cell proliferation and differentiation. Oncogenic activation of SMO leads to overactivation of GLI transcription and numerous other pathways, revealing the essential role of GPCRs in regulating a multiplicity of downstream effectors. Our study provides a detailed analysis of the cellular and transcriptional effects of dysregulated GPCR signaling in the skin stem cell compartment. Despite the hair follicle characteristics of BCC, experimental data in mice indicates that hair follicle stem cell populations might not be the main origin of tumors ^18,19,37^. Here, we demonstrate that BCC-like cells cluster in two distinct populations with markers for touch-dome and isthmus stem cell-like cells. Interestingly, we find that while bulge *Lgr5*+ stem cells are not able to generate BCC-like tumors, *Lrig1*+ isthmus stem cells have limited capacity to generate invasive BCC lesions following PKA inactivation. These results suggest that the main population of invasive BCC-like lesions originates outside of the hair follicle, mainly from touch-dome stem cell-like cells. Indeed, touch-dome *Gli*+ cells express broad interfollicular and hair follicle stem cell characteristics, similar to BCCA cells, and have been proposed to be the origin of BCC lesions in previous studies ^18,19^.

BCC has been classically characterized by genomic alterations in Hedgehog signaling members, although BCCs present numerous genetic changes in other growth-promoting and differentiation-suppressing mechanisms ^3,38,39^. In addition, some patients with high-frequency BCCs do not show germline mutations in the Hedgehog pathway ^39,40^. This evidence indicates that additional alterations could drive BCC formation in humans. Indeed, our results show disruptive mutations in Gαs and PKA in human tumors, suggesting that they could contribute to human BCC. However, the low burden of Gαs and PKA genomic alterations indicates that additional genetic and signaling mechanisms might exist in BCC to inactivate Gαs and PKA. Bazex-Dupré-Christol syndrome, a rare disease that presents with multiple BCCs, has been linked to amplifications in the *ARHGAP36* gene ^41^. ARHGAP36 can antagonize PKA and activate Hedgehog signaling ^42^, establishing ARHGAP36 amplification as an alternative pathway to achieve PKA inactivation in BCC. Further research is needed to unveil additional mechanisms by which Gαs pathway inactivation contributes to human BCC and other tumors.

PKA activation has been proposed as a therapy to block oncogenic Hedgehog signaling, although this has not been studied in detail in skin BCC. Here, we find that blockage of cAMP degradation by PDE inhibitors does not affect the growth of SmoM2-induced tumors, but activation of the Gαs-coupled ADORA2B can lead to reduced tumor burden and proliferation. We also find potential differences between Gαs-coupled GPCR agonists, indicating that general activation of the Gαs pathway might not be a viable strategy to block oncogenic Hedgehog signaling. It has been demonstrated that Gαs-coupled GPCR activation is involved in T-cell exhaustion ^43^. Interestingly, ADORA2B is not expressed in T-cells, while adrenergic receptors are enriched in exhausted T-cells ^43^. Our results show that targeting Gαs-coupled GPCRs specifically expressed by tumor cells can enhance cAMP-mediated therapy. It is worth noting that the SmoM2 mouse model represents SMO-inhibitor-resistant BCC ^10^, indicating that targeting ADORA2B could be an alternative option for drug-resistant tumors.

The mouse models we chose for our study were based on the main objective to characterize the relationship between Gαs pathway inactivation and canonical Hedgehog signaling. However, these BCC mouse models present some limitations. BCC is one of the human cancers with the highest mutational burden ^39,44,45^, a fact that is not recapitulated in our single-oncogene mouse BCC-like lesions. In addition, SmoM2 tumors are sometimes classified as basaloid follicular hamartomas rather than BCC ^46,47^. Hamartomas are a type of benign skin lesion with lower Hedgehog activation than BCC ^46,48^. Despite this distinction, SmoM2 mice carry a mutation present in human BCC patients and recapitulate BCC development ^49^. SmoM2 mice are extensively used as a model to study BCC, oncogenic Hedgehog signaling, and SMO-inhibitor-resistant BCC.

Our study provides a valuable resource to understand the developmental and pathological roles of Gαs and Hedgehog signaling pathway crosstalk. The findings also reveal that oncogenic GPCR activation, such as SMO-driven signaling imbalances, can be counteracted by activating complementary pathways, restoring GPCR signaling equilibrium and normal physiological processes essential for tissue homeostasis. Further research is needed to identify and expand the repertoire of signaling modulators capable of reestablishing normal cycles of stem cell proliferation and differentiation, ultimately restoring normal tissue function.

## MATERIALS AND METHODS

### Mice

All mouse studies were carried out according to approved protocols from the National Institutes of Health Intramural Animal Care and Use Committee of the National Cancer Institute (Bethesda, MD), in compliance with the Guide for the Care and Use of Laboratory Animals. The following mouse lines were obtained from the Jackson Laboratory: LSL-SmoM2 (stock 005130), LSL-rtTA (stock 005670), K14CreERTM (stock 005107), *GliLz* reporter mice (stock 008211), K14Cre (stock 018964), *Smo*-floxed mice (stock 004526). *Gnas*-Floxed mice were provided by Lee Weinstein (NIDDK, NIH) ^11^. *Gpr161*-floxed mice ^30^ and *Gpr124*-floxed mice ^28^ have been described before. tetO-PKI transgenic mice were generated with the assistance from the CCR Transgenics Facility. A codon-optimized sequence for the 1–24 amino acids from human PKIA downstream of GFP was cloned into a vector with a seven tet-responsive element (TRE) and flanked by the chicken h-globin gene insulator (HS4) as described before ^50^. Founders were identified for the transgene by screening genomic DNA from tail biopsies using a PCR reaction with the following primers: forward sequence 5′-GGGGAAGGGGATGCTACCTA-3′, reverse sequence 5′-TCGAATTTCACTTCTGCCCGA-3′, band approximately 230 bp. PCR reactions were performed as follow: 95 °C for 2 min, followed by 30 cycles of 95 °C for 30 s, 62 °C for 45 s, and 72 °C for 45 s, and a final cycle with 2 min of extension at 72 °C. All experiments were conducted using littermate controls and both male and female mice were used. To identify knockout bands in *Gpr124* and *Gpr161*-floxed animals, DNA samples from tail were used for PCR with the following cycles, *Gpr124*: 94°C for 2 min, followed by 32 cycles of 94°C for 30sec, 60°C for 30sec, and 70°C for 1 min, and a final cycle with 5min at 70°C; *Gpr161*: 94°C for 2 min, followed by a touchdown protocol of 10 cycles of 94°C for 20sec, 65°C (-0.5°C per cycle decrease) for 15sec, and 68°C for 10sec, followed by 30 cycles of 94°C for 15sec, 60°C for 15sec, and 72°C for 80 sec, and a final cycle with 2min at 72°C. Primer used were: Gpr124-F, 5’-TGGTTGCCTTACAAGTCTGAGC-3’; Gpr124-R1, 5’-CTGATACCCTGCCAGGCG-3’; Gpr124-R2, 5’-CGGTTGTCGCCTGGACTC-3’; Gpr161_2R, 5’-TGAACTGATGGCGAGCTCAGACC-3’; Gpr161_3F, 5’-CAAGATGGATTCGCAGTAGCTTGG-3’; Gpr161_4R, 5’-ATGGGGTACACCATTGGATACAGG-3’.

For CreERTM activation, mice were injected intraperitoneally with one dose of 100mg/kg tamoxifen (Sigma T5648) between 8 and 12 weeks of age. PKI expression was induced by doxycycline treatment immediately following tamoxifen administration. Doxycycline was administered in food grain-based pellets (Bio-Serv, Flemington, NJ) at 6g/kg. For drug treatments, SmoM2 mice were treated with tamoxifen and then randomized into each treatment group by utilizing a random number generator. For individual PDE inhibitor treatment, rolipram (Cayman Chemicals 10011132) and BRL50481 (Cayman Chemicals 16899) were injected intraperitoneally at 10mg/kg/day for 29 days, 8 days after tamoxifen induction. For combined treatment, rolipram and BRL50481 were injected together at a dose of 10mg/kg/day each for 8 days, 40 days after tamoxifen induction. Control mice were injected with vehicle. For topical drug treatments, isoproterenol (Sigma I5627) 0.25mg/site/day, and BAY 60-6583 (Tocris 4472) 10μg/site/day, in DMSO, were applied to the ear of mice for 10 days. These doses have been used before successfully for topical skin treatment ^51,52^. DMSO alone was applied to ears of control mice.

### DNA constructs

Genes were cloned using gBlocks Gene Fragments (Integrated DNA Technologies) coding for the corresponding amino acid sequence into a pCEFL vector. *GPR161* was cloned utilizing a codon-optimized version based on UniProt Q8N6U8-1. GPR161-V129E mutation was introduced utilizing QuikChange II Site-directed mutagenesis kit (Agilent). For inducible lentivirus production, GPR161 constructs were transferred into the pInducer20 vector (Addgene plasmid #44012). *GNAS* was cloned using as a base Gs long EE-tag (internal) from Missouri S&T cDNA Resource Center (Catalogue Number: GNA0SLEI00), codon optimized for DNA synthesis and individual mutations were added to each gBlock construct. PRKACA was cloned based on NM_002730.4 with an added C-terminal HA tag, codon optimized for DNA synthesis and individual mutations were added to each gBlock construct. pGL4.29[luc2P/CRE/Hygro] (CRE-Luc) was purchased from Promega. SV40 LargeT lentivirus vector was from Addgene (LV-EF1a-SV40-LargeT-Antigen-P2A-Hygro, #170255).

### Cell culture and transfections

All cells were cultured at 37°C in the presence of 5% CO2. Lenti-X 293T cells for lentiviral production were obtained from Takara Bio (632180). HEK293 cells were obtained from AddexBio (T0011001). These cells were cultured in DMEM (Sigma D5796) containing 10% fetal bovine serum (Sigma F4135) and antibiotic/antimycotic solution (Sigma A5955). N/TERT2G keratinocyte cell line was cultured in EpiLife media (Gibco MEPI500CA) with Human Keratinocyte Growth Supplement (HKGS, Gibco S0015) as previously described ^50^. Transfections were performed with Lipofectamine 3000 (Invitrogen) according to the manufacturer’s instructions. To generate a SmoM2 BCC cell line, tumor cells were isolated from the tail skin of SmoM2 mice 40 days post-tamoxifen injection. Tail skin was removed from mice, cleaned using 10% iodine in PBS and washed with sterile PBS several times. Each tail was cut into pieces and then incubated overnight at 4°C in 2.5 parts of 0.25% trypsin (Gibco 15050-057) and 1 part of 5U/mL dispase (Stem Cell Technologies 07913). The next day, the epidermis was scraped from the dermis and minced in 0.25% trypsin with EDTA (Gibco 25200056). Trypsin was neutralized in DMEM with 10% FBS and cells were strained in a 100μm cell strainer followed by a 40μm cell strainer. After centrifugation at 500G for 8 minutes cells were plated on Culturex BME2 (3533-005-02) coated plates in EpiLife + HKGS + 10μM Y-27632 (Cayman 10005583) + 10ng/mL mouse EGF (Peprotech 315-09). After two passages, cells were transduced with SV40 LargeT-expressing lentiviruses and selected with 200 μg/mL hygromycin for two days (Gibco 10687010). After six passages, one million cells were injected subcutaneously into the flank of NSG mice to select for oncogenic cells. Eighty-four days later, the xenograft tumor was harvested, trimmed, incubated in 10% iodine solution in PBS and washed several times with sterile PBS. Then the tumor was minced and incubated in a solution of dispase (1 U/mL) and Type I collagenase (200 U/mL, Worthington Biomedical LS004214) at 37°C for two hours. The tumor was then strained in a 100μm cell strainer, and the strainer was washed with DMEM + 10% FBS. After centrifugation at 500G for 8 minutes, the tumor was plated in EpiLife + Human Keratinocyte Growth Supplement + 10μM Y-27632 + 10ng/mL mouse EGF. After two passages, EpiLife medium was switched to high glucose DMEM + 10% FBS + 10μM Y-27632 + 10ng/mL mouse EGF. After four passages, cells were sorted with FACS (BD FACS Aria III machine) for high YFP expression. The sorted cells were injected into the flank of an NSG mouse, and the xenograft tumor was harvested, lysed, and plated for cell culture in high glucose DMEM medium + 10% FBS + 10μM Y-27632 + 10ng/mL mouse EGF. This final cell stock was used for experiments. Population doubling was performed as previously described ^53^. SmoM2 cells were validated by STR profiling (ATCC) with the following results: Locus 18-3:17; 4-2:18.3; 6-7:15; 19-2: 12, 13; 1-2: 13, 19; 7-1: 27.2, 30; 1-1: 17; 3-2:12, 14; 8-1: 16, 17; 2-1: 9; 15-3: 20.3, 21.3, 22.3, 6-4: 16.3, 18; 11-2: 16; 17-2: 13; 12-1: 20; 5-5: 17; X-1: 27; 13-1: 15, 17; human and/or African green monkey markers were not detected. All cells were routinely tested for mycoplasma by PCR.

### Luciferase Assays

Cells in 24-well plates were co-transfected with CRE-Luc (105 ng/cm2) plus the indicated DNA constructs (99 ng/cm2), and a Renilla Luciferase Vector (21 ng/cm2). Luciferase activity was measured 24 hours after transfection using a Dual-Glo Luciferase Assay Kit (Promega E2940) and a Microtiter plate luminometer (SpectraMax iD3, Molecular Devices LLC). Firefly luciferase activity was normalized to renilla luciferase in every sample.

### Immunofluorescence and staining

For staining of cilia in cells, SmoM2 cells were seeded on coverslips in 24-well plates in complete medium (DMEM + 10% FBS + 10 μM Y-27632 + 10 ng/mL mouse EGF). Cells were serum starved (DMEM with no supplements) overnight and then fixed in methanol at -20°C for 1 hour. Cells were washed with block solution (PBS + 5% FBS + 0.2% Tween 20) and stained with primary antibody (1:100 ARL13B, BiCell Scientific 90413) overnight at 4°C. After washing with block solution, coverslips were stained with secondary antibody (1:200 Donkey anti-Rat, Invitrogen SA5-10027) for 1 hour. For skin whole mount staining, tail skin was isolated and incubated in 5mM EDTA for 4 hours at 37°C. Using tweezers, the epidermis was then separated from the dermis. The epidermal sheets were fixed in 3% paraformaldehyde for one hour and then submerged in blocking buffer (PBS + 10%FBS + 0.3% Triton-X + 0.02% sodium azide) for one hour. Next, primary antibody was added in blocking buffer (1:500 Keratin 5-Biolegend 905901, 1:400 Keratin 17-Cell Signaling 12509, and 1:500 Acetyl-α-Tubulin-Cell Signaling 5335) overnight at 4°C in a humidity chamber. After washing three times for 30 minutes in washing buffer (PBS + 1%FBS + 0.2% Triton X-100 + 0.02% sodium azide), 1:400 secondary antibody was added in blocking buffer (Alexa Fluor Donkey Anti-Rabbit 555 Invitrogen A31572 and Alexa Fluor Goat Anti-Chicken 633 Invitrogen A21103). After three washes in washing buffer, 10 μg/mL Hoechst 33342 (Molecular Probes H-3570) was added in PBS for 10 minutes. Immunofluorescence analysis of mouse skin was performed on tissue sections embedded in paraffin as previously described ^50^ with PCNA antibody (1:400 Cell Signaling 13110S). The stained epidermis was then mounted with FluorSave Reagent (EMD Millipore 345789) on charged microscope slides (Globe Scientific 1358G) with No 1.5 coverglass (Thermofisher 3422). Whole mount and paraffin embedding methods quench transgene YFP and GFP signal. Slides were imaged on a Leica SP8 confocal microscope with LASX software (high-contrast Plan-Apochromat 20x oil CS2 1.30 NA objective; Leica Microsystems) or with a Keyence BZ-X700 with BZX software (CFI Plan Apo λ20x NA 0.75 objective, Nikon). Whole mount beta-galactosidase staining was performed as previously described ^7^. For histological analysis tail and ear skin or tumors were fixed in Z-Fix (Anatech Ltd 174) overnight and then stored in 70% ethanol. Tissues were embedded in paraffin and 3-μm sections were stained with H&E. Stained H&E slides were scanned at 40x using an Aperio CS Scanscope (Leica Microsystems) or a NanoZoomer Digital Slide Scanner (Hamamatsu). Tumor burden level was quantified in the ear H&E sections by HALO image analysis platform (Indica Labs). The total epidermal area was calculated and then divided by length of the tissue sections. Final images were assembled in AdobeIllustrator 29.2.

### Immunoblotting

Cells were lysed by sonication in lysis buffer: 50mM Tris-HCl pH 8.0, 150mM NaCl, 1mM EDTA, 1% NP40, 0.2% sodium deoxycholate, 0.1% SDS, 1mM dithiothreitol (Sigma D9779), protease inhibitor (cOmplete ULTRA tablets Roche 766534), and phosphatase inhibitor (PhosSTOP tablets Roche 04906837001). Protein quantification was performed with BioRad Protein Assay kit. An equal amount of protein was loaded onto SDS-polyacrylamide gels for electrophoresis and transferred to PVDF membranes. Antibodies used were: GAPDH (1:2000, Cell Signaling 5174), HA Tag (1:1000, Cell Signaling 3724), Gli3 (1:300, R&D Systems AF3690), SMO (1:50, Santa Cruz sc-166685), E-Cadherin (1:1000, Cell Signaling 3195), Keratin 17 (1:2000, Cell Signaling 12509), EE Tag (1:300, Biolegend 901801), phospho-PKA substrates (1:1000, Cell Signaling 9624), phospho-CREB Ser133 (1:1000, Cell Signaling 9198), and total CREB (1:1000, Cell Signaling 9197). Secondary horseradish peroxidase-conjugated antibodies used were Pierce peroxidase goat anti-mouse IgG (HþL) (Thermo Fisher Scientific 31432; 1:4,000), Proteintech HRP-Goat anti-rabbit (RGAR001; 1:20,000), R&D anti-goat (HAF109; 1:1000). Bands were detected using a ChemiDoc Imaging System (Bio-Rad Laboratories) with Clarity Western ECL Blotting Substrates (Bio-Rad Laboratories) according to the manufacturer’s instructions. Blot images were processed using ImageLab software, version 5.2.1 (Bio-Rad Laboratories). Final images were assembled in AdobeIllustrator 29.2.

### Gene expression analysis and quantitative PCR

Gene expression analysis and single-cell sequencing was performed from isolated tail keratinocytes as described in the Cell culture section. For single-cell RNA sequencing, cells were collected in EpiLife media after centrifugation and processed. For bulk RNA sequencing, cells were lysed in RLT Lysis buffer from the RNeasy Plus Mini Kit. For bulk RNAseq, keratinocytes were isolated from SmoM2 mice 29 days, *Gnas*-eKO mice 42 days, and PKI mice 57 days, post-tumor induction. For single-cell RNAseq, keratinocytes were isolated from all mice 28 days post-tumor induction. For *Gpr161*-eKO mice RNA sequencing, mice were harvested at four months of age. For bulk RNAseq and quantitative PCR, isolated keratinocyte RNA was purified using RNeasy Plus Mini Kit (Qiagen 74134) according to the manufacturer’s instructions. Cells were lysed using the Precellys lysing kit (Bertin Instruments). To synthesize cDNA, one microgram of RNA was used with the SuperScript IV VILO kit (Invitrogen). The Taqman Fast Advanced Master Mix (ThermoFisher 4444557) was used with Taqman probes for quantitative PCR. Targets are as follows: *Smo* (Mm011672710_m1), *Ptch1* (Mm00436026_m1), *Gli1* (Mm00494654_m1), *Gli2* (Mm01293117_m1), and *Rn18s* (Mm03928990_g1). Values for Taqman probes were normalized using *Rn18s* expression. For expression of *Gpr161*, PowerSybr Green PCR Master Mix (ThermoFisher 4367659) was used with a PrimeTime qPCR primer for m*Gpr161* (IDT Mm.PT.58.28865637) and primers for housekeeping gene m*Rps14* for normalization (*Rps14*-F, 5’-GACCAAGACCCCTGGACCT-3’ and *Rps14*-R, 5’-CCCCTTTTCTTCGAGTGCTA -3’). Samples were analyzed using QuantStudio 5 Real-Time PCR System (Thermo Fisher Scientific).

For bulk RNAseq, mRNA expression profiling was performed by the Center for Cancer Research Sequencing Facility, National Institutes of Health. Reads were trimmed for adapters and low-quality bases using Trimmomatic software and aligned with the reference genome mouse-mm10. Transcripts were annotated using STAR. Gene counts for genes with ≥5 reads were normalized to the trimmed mean of M values using Partek Flow software, version 12 (Partek). Normalized counts were used for differential analysis using Partek Flow GSA algorithm (Partek). Kegg pathway enrichment was performed using Partek. Upstream regulators analysis was performed using Ingenuity Pathway Analysis (Ingenuity Systems, www.ingenuity.com). For Single-cell RNA-seq from our BCC mouse models, tail keratinocytes were isolated from two mice of each genotype, 3 weeks following tamoxifen/doxycycline induction. Single-cell isolation and data processing was performed by the CCR Single Cell Analysis Facility. 10X Genomics Chromium Next GEM Single Cell 3’v3.1 (Dual Index) was used with a single capture lane per sample. Sequencing was performed by the Center for Cancer Research Sequencing Facility, National Institutes of Health. Base calling was performed using RTA 3.4.4, demultiplexing was performed using cellranger v7.0.1 (Bcl2fastq 2.20.0), and alignment was performed using cellranger v7.0.1 (STAR 2.7.2a). Sequenced reads were aligned to the 10x Genomics provided mouse reference sequence (refdata-gex-mm10-2020-A). Single-cell counts were imported and analyzed using Partek Flow software, version 12 (Partek). Cells were filtered by quality measurements and counts were normalized using counts per million. Normalized counts for each cell were used for unsupervised clustering using principal component analysis (PCA), followed by Harmony batch removal tool ^54^ and graph-based clustering to identify cell groups. Biomarkers for each cell population were calculated, and cells were classified by expression of known markers for each cell type ^15–18^. Only basal cell populations were selected for further analysis. Clustering of groups was made by creating pseudobulk data in Partek Flow by pooling single cells from each corresponding group using sum of raw counts as aggregation method.

### BCC mutation analysis

Human BCC mutations were obtained from ^25^. Only mutations present in *GNASL* (NM_000516) and *GNASS* (NM_080426) were considered since these are the only *GNAS* variants expressed in keratinocytes. Only mutations in coding sequences were further analyzed. We predicted the impact of mutations through several algorithms available via the Ensembl Variant Effect Predictor (VEP) ^55^, such as sift, polyphen2, mutation assessor, cadd, alphamissense, primateai, spliceai, mutation taster, dbscsnv and mutpred. The effect of mutationss were further analyzed through the Atlantis webserver (https://atlantis.bioinfolab.sns.it), which provides a framework for understanding the functional role of specific residues in protein structures. In particular, we used pre-calculated 3D contacts to characterize the residue at the interfaces with the mutated amino acids.

### In-vivo microscopy and Imaris

The non-invasive procedure for skin imaging was performed as previously described ^56^. The imaging was performed in the ear of 3 mice on days 3 and 17 after induction by using an inverted TCS SP8 Dive Spectral Microscope (Leica) equipped with a Mai-Tai tunable laser (Spectral Physics), 2 sepctral detectors (HyD-RLD, Leica) and a 37 °C preheated 40X objective (NA 1.10, HC PL IRAPO, Leica). The specimens were excited at 910 nm. Collagen I (Second Harmonic Generation) and GFP were detected respectively at 451 – 461 nm and 504 – 526 nm. The tile mode of the samples was collected by bidirectional line scanning at 400 Hz (320 x 320 pixel; 12 bits per pixel) on XY, and 40-50 Z-stacks (2 um step size) using LAS X Navigator software. All the images were then stitched together and stored as LIF files for further processing using Imaris (Bitplane). For image analysis, Imaris 9.2.1 (Bitplane AG) software was used. The surfaces algorithm was applied to GFP-PKI tumors to calculate tumor volume. The measurements point algorithm was used to measure the distance between tumors and hair follicles.

### Statistical analysis

The number of independent experiments and statistical tests are indicated in each figure legend. Statistical analyses were carried out using Prism 7 (GraphPad Software). Asterisks denote statistical significance.

### Data availability

RNA-sequencing datasets generated for this study are available in the Gene Expression Omnibus database under accession codes GSE290008 (Bulk RNAseq BCC models), GSE290004 (*Gpr161*eKO), and GSE290180 (single-cell RNAseq BCC models).

## AUTHOR CONTRIBUTIONS

Conceptualization RIB; Data curation SK, BB, KL, NSP, YN, NDOR, FR, RIB; Formal analysis SK, BB, KL, NSP, YN, NDOR, KYS, FR, RIB; Funding acquisition RIB; Investigation SK, BB, KL, NSP, YN, NDOR, KYS, FR, RIB; Project administration SK, RIB; Resources SM, BSC, RW, FR, RIB; Writing – original draft SK, RIB; Writing – review & editing SK, BB, KL, NSP, YN, NDOR, SM, BSC, KYS, RW, FR, RIB.

## ACKNOWLEDGMENTS

This research was supported by the Intramural Research Program of the National Institutes of Health, National Cancer Institute, Center for Cancer Research (ZIA BC 011763). S.M. was supported by grant R35GM144136. This work used the computational resources of the National Institutes of Health High-Performance Computing Biowulf Cluster. Analysis and management of H&E images were supported by the National Cancer Institute HALO Image Analysis Resource. The authors thank the members of the CCR Sequencing Facility at Frederick National Laboratory for Cancer Research and the CCR Single Cell Analysis Facility for their help during sample preparation, sequencing, and data processing.

---

**Figure S1.**
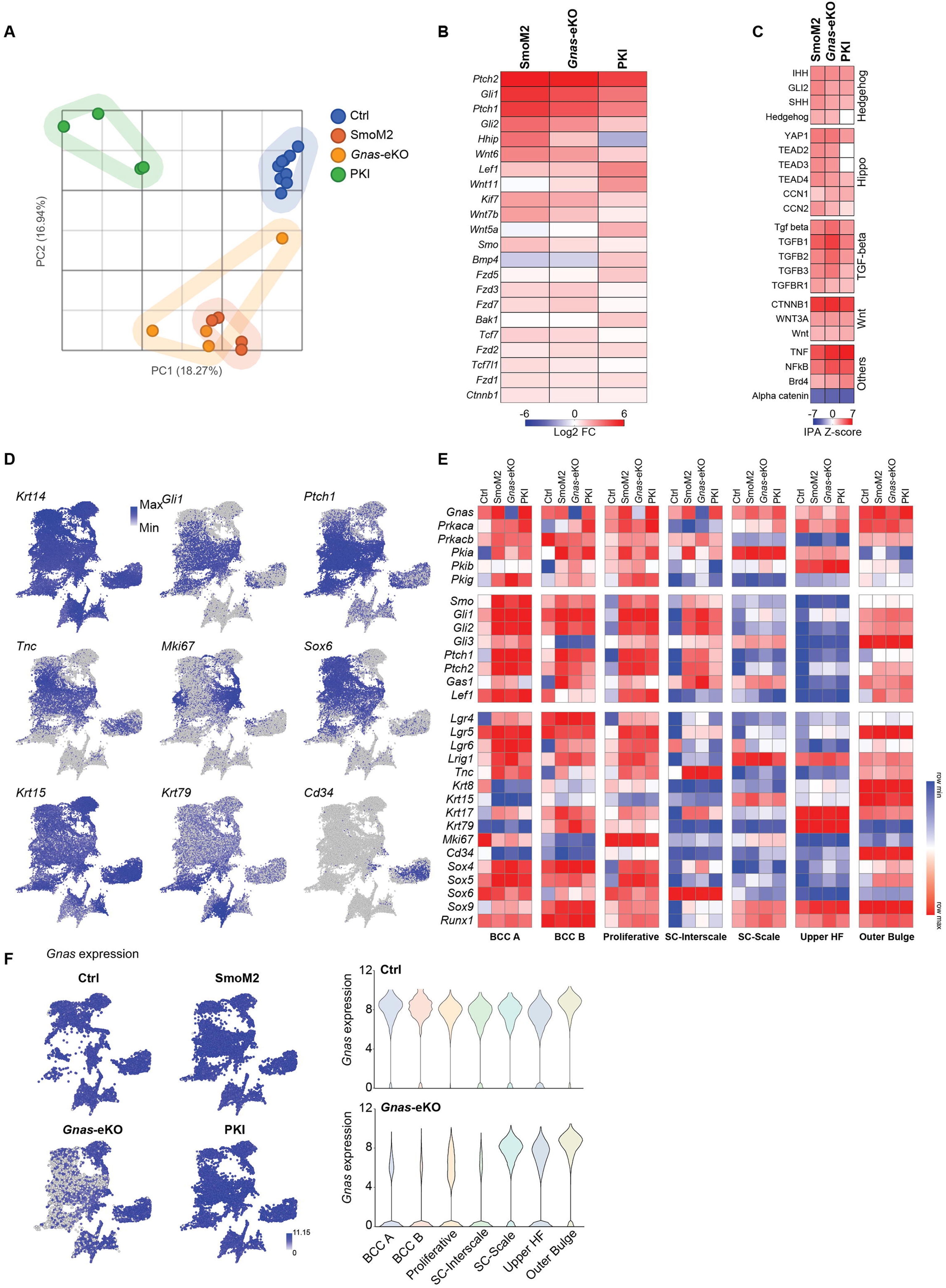
**A-** PCA plot showing clustering of the different mice by their gene expression profile in bulk mRNA-seq. **B-** Heat map from bulk mRNA-sequencing data comparing the fold change (log2FC) increased in BCC-related genes in the indicated mice. **C-** Functional analysis of pathways differentially regulated in BCC mouse models with respect to control mice. Generated using profile bulk mRNA-seq data with Ingenuity Pathway Analysis software (IPA, Ingenuity Systems). **D-** Expression of cluster-defining genes projected into UMAP of cell clusters for all mice (Fig. 2C). **E-** Relative expression of the indicated genes in each cell compartment for each mouse genotype from single-cell RNAseq data. **F-** Left: expression of *Gnas* projected into a UMAP of each mouse model. Right: violin plot depicting the expression of *Gnas* for individual cells in each compartment in control (Ctrl) and *Gnas*-eKO mice.

**Figure S2.**
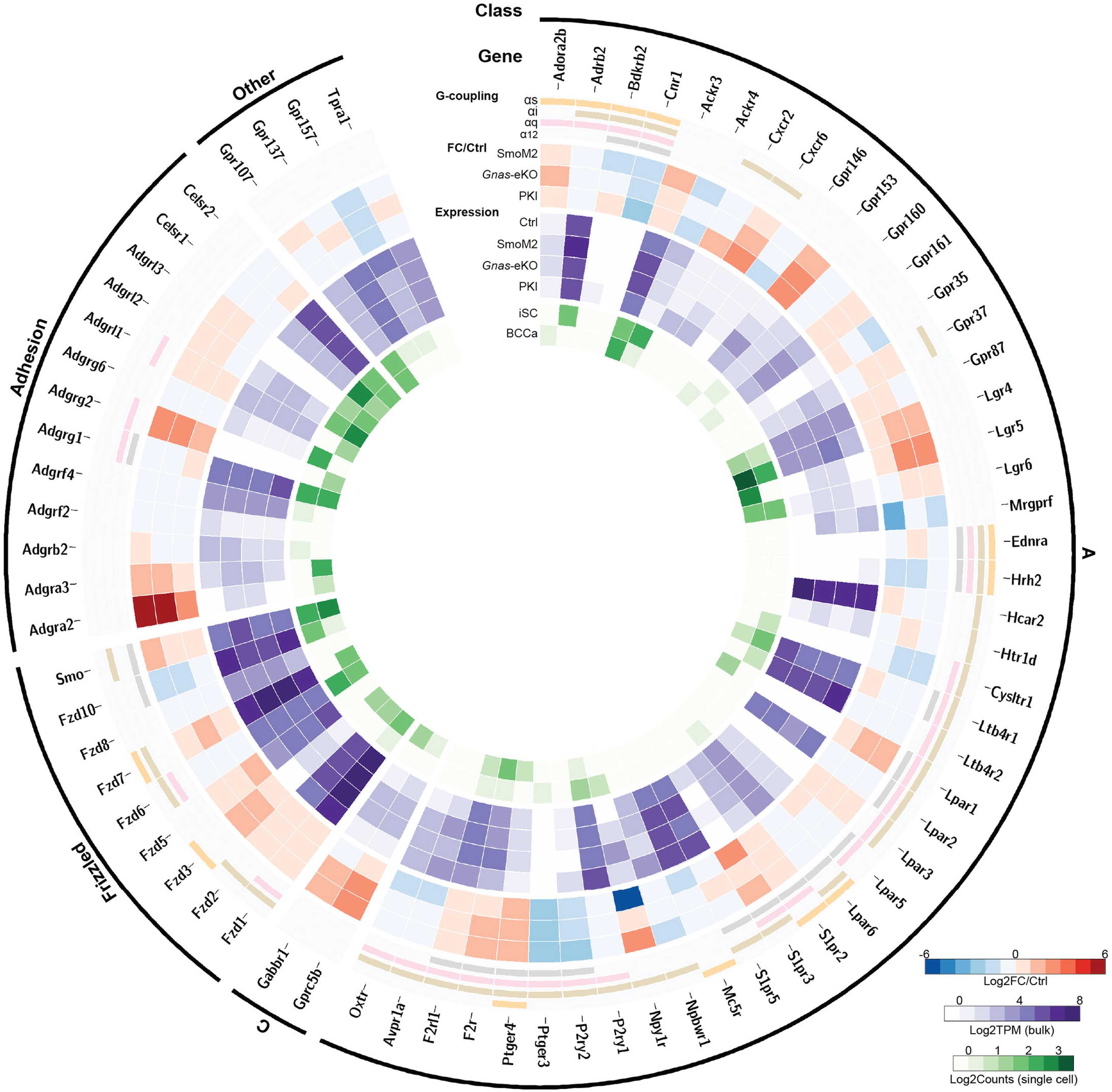
Amplified version of Figure 5A. Circos plot representing BCC GPCR expression in mice, showing class, gene name, Gα coupling, fold change (FC) and expression level (Log2TPM) in bulk RNAseq, level of expression in single-cell RNA sequencing (Log2Counts) in interscale (iSC) and BCCA clusters.

**Figure S3.**
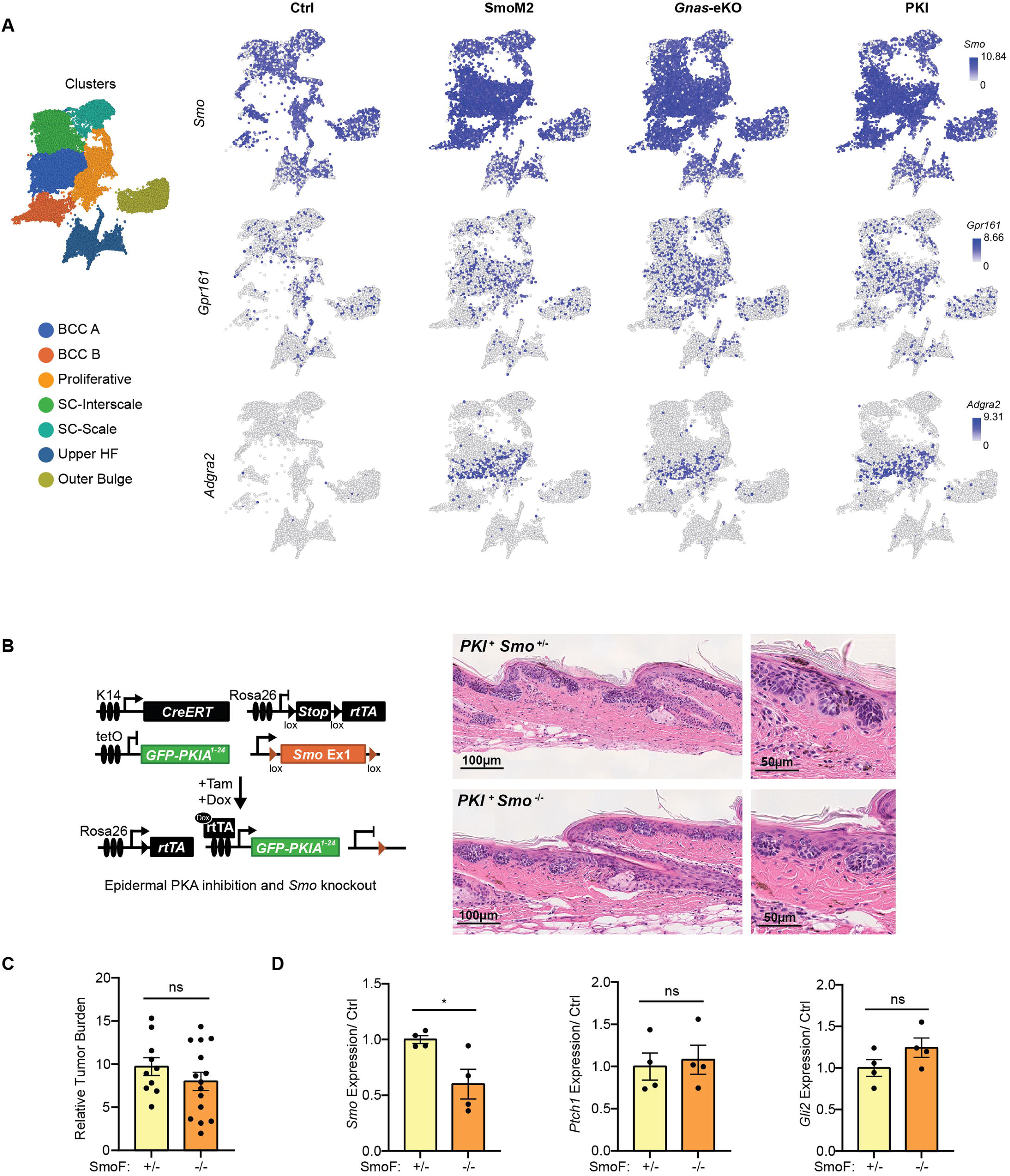
**A-** Expression of the indicated GPCRs projected into a UMAP of each mouse model. UMAP of cell clusters for all mice (Fig. 2C) and names of cell clusters is included for reference. **B-** Schematic representation of the mouse model used to target SMO deletion and PKI expression to the skin. Images on the right show representative H&E staining of tail skin from the indicated mice 34 days after induction. + indicates transgene is present, -/+ indicates heterozygous knockout and -/- indicates homozygous knockout. **C-** Quantification of tumor burden per mouse in H&E tail skin stainings in PKI^+^ mice with *Smo* heterozygous (+/-) or homozygous (-/-) knockout. Graphs show mean ± SEM. N= 10 SmoF +/- mice, N= 14 SmoF -/- mice, t-test. **D-** qRT-PCR analysis of mRNA expression of indicated markers in tail epidermal keratinocytes isolated from PKI^+^ mice with *Smo* heterozygous (+/-) or homozygous (-/-) knockout 34 days after induction. Graphs show mean ± SEM. N= 4 SmoF +/- mice, N= 4 SmoF -/- mice, t-test. SmoF +/- are used as control (Ctrl).

**Figure S4.**
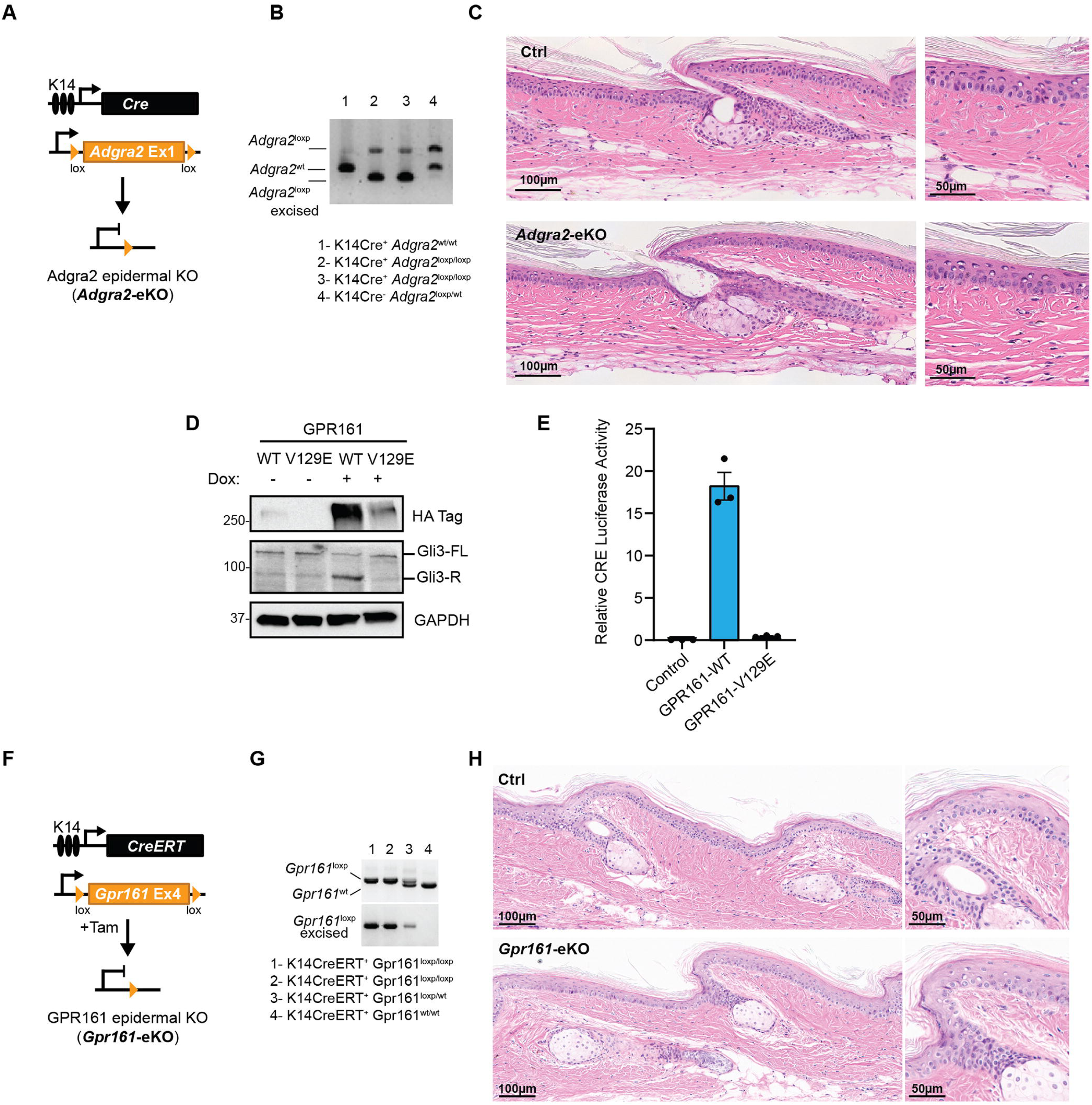
**A-** Schematic representation of the mouse model used to target *Adgra2* deletion to the skin. **B-** Validation of effective *Adgra2* deletion by PCR in the indicated mice. **C-** Representative H&E staining of tail skin from the indicated mice 26 weeks after induction. **D-** Western blot analysis of expression of Gli3-Repressor and expression of doxycycline (Dox)-inducible GPR161-WT and inactivating mutation GPR161-V129E constructs in NTERT human keratinocytes. Molecular weight markers (kDa) are indicated on the left. **E-** Transcriptional activity of CREB measured by CRE-luciferase assay in HEK293 cells transfected with the indicated constructs. Graphs show mean ± SEM. N=3. **F-** Schematic representation of the mouse model used to target *Gpr161* deletion to the skin. **G-** Validation of effective *Gpr161* deletion by PCR in the indicated mice. **H-** Representative H&E staining of tail skin from the indicated mice 90 days after induction.

**Fig S5.**
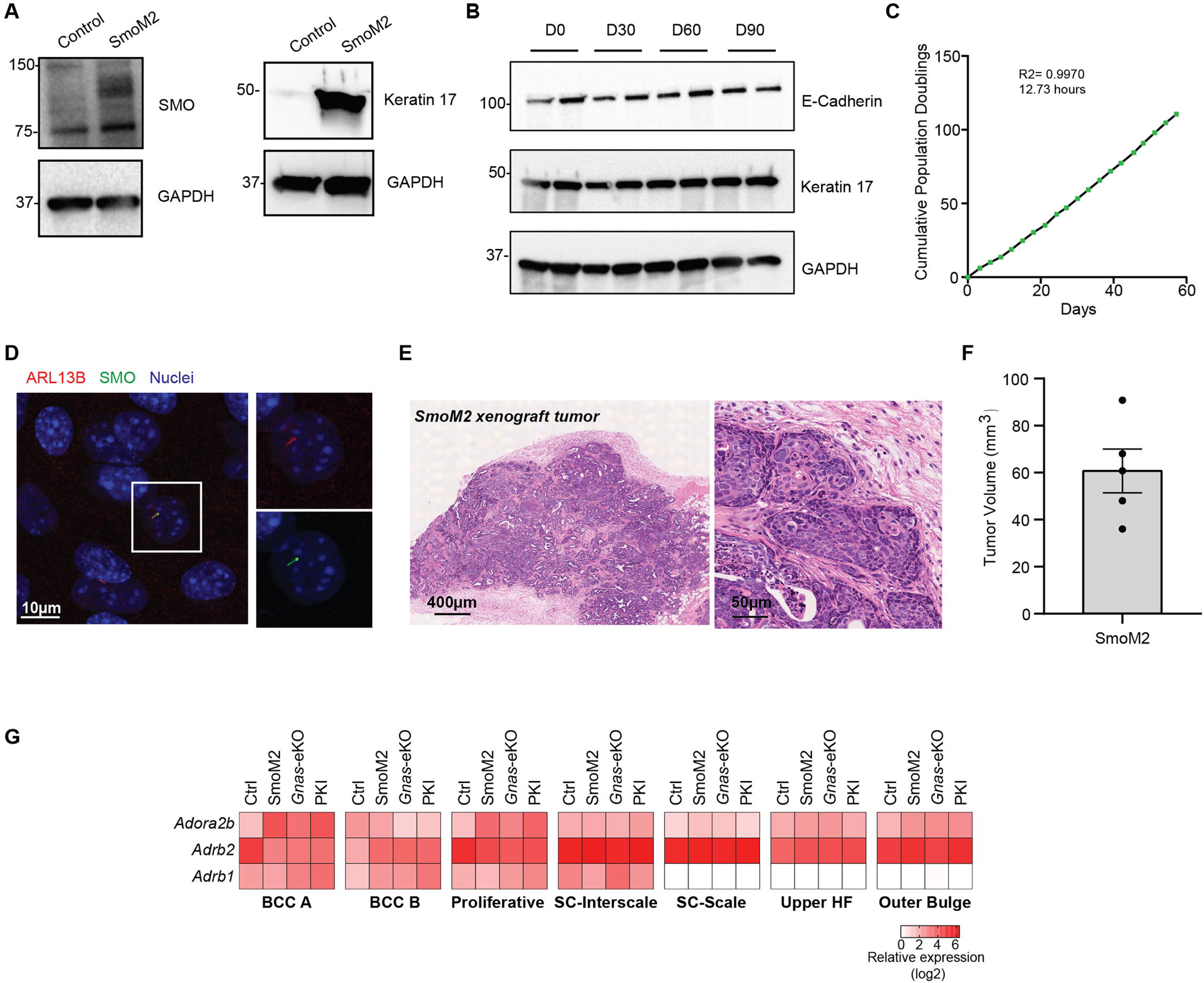
**A-** Western blot analysis of SMO and Keratin 17 expression in a SmoM2 cell line generated from BCC tail tumors from SmoM2 mice. Molecular weight markers (kDa) are indicated on the left. **B-** Western blot analysis of E-Cadherin and Keratin 17 expression in SmoM2 cell culture model after 30, 60, and 90 days (D) in culture. Molecular weight markers (kDa) are indicated on the left. **C-** Population doubling graph of SmoM2 cells over 60 days with best line fit regression curve and average doubling time. **D-** IF of SmoM2 cells with magnified field showing the cilia marker ARL13B (red) and ciliary localization of SMO (green). SmoM2 mice and cells express SMO fused to YFP protein. **E and F-** Representative images from H&E staining and quantification of tumor volume of SmoM2 xenograft tumors from subcutaneous injection of SmoM2 keratinocytes in the flank of NOD/SCID mouse, 35 days after injection. **G-** Heat map from single cell RNAseq data from the indicated cell groups comparing the relative expression of the Gαs-coupled GPCRs among mouse models.

## REFERENCES

1. Pedro, M.P., Lund, K., and Iglesias-Bartolome, R. (2020). The landscape of GPCR signaling in the regulation of epidermal stem cell fate and skin homeostasis. Stem Cells 38, 1520–1531. 10.1002/stem.3273.

2. Kong, J.H., Siebold, C., and Rohatgi, R. (2019). Biochemical mechanisms of vertebrate hedgehog signaling. Development 146. 10.1242/dev.166892.

3. Epstein, E.H. (2008). Basal cell carcinomas: attack of the hedgehog. Nature reviews. Cancer 8, 743–754. 10.1038/nrc2503.

4. Rogers, H.W., Weinstock, M.A., Feldman, S.R., and Coldiron, B.M. (2015). Incidence Estimate of Nonmelanoma Skin Cancer (Keratinocyte Carcinomas) in the U.S. Population, 2012. JAMA Dermatol 151, 1081–1086. 10.1001/jamadermatol.2015.1187.

5. Sekulic, A., and Von Hoff, D. (2016). Hedgehog Pathway Inhibition. Cell 164, 831. 10.1016/j.cell.2016.02.021.

6. Peris, K., Fargnoli, M.C., Kaufmann, R., Arenberger, P., Bastholt, L., Seguin, N.B., Bataille, V., Brochez, L., Del Marmol, V., Dummer, R., et al. (2023). European consensus-based interdisciplinary guideline for diagnosis and treatment of basal cell carcinoma-update 2023. Eur J Cancer 192, 113254. 10.1016/j.ejca.2023.113254.

7. Iglesias-Bartolome, R., Torres, D., Marone, R., Feng, X., Martin, D., Simaan, M., Chen, M., Weinstein, L.S., Taylor, S.S., Molinolo, A.A., and Gutkind, J.S. (2015). Inactivation of a Galpha(s)-PKA tumour suppressor pathway in skin stem cells initiates basal-cell carcinogenesis. Nature cell biology 17, 793–803. 10.1038/ncb3164.

8. Vasioukhin, V., Degenstein, L., Wise, B., and Fuchs, E. (1999). The magical touch: genome targeting in epidermal stem cells induced by tamoxifen application to mouse skin. Proceedings of the National Academy of Sciences of the United States of America 96, 8551–8556. 10.1073/pnas.96.15.8551.

9. Mao, J., Ligon, K.L., Rakhlin, E.Y., Thayer, S.P., Bronson, R.T., Rowitch, D., and McMahon, A.P. (2006). A novel somatic mouse model to survey tumorigenic potential applied to the Hedgehog pathway. Cancer research 66, 10171–10178. 10.1158/0008-5472.CAN-06-0657.

10. Atwood, S.X., Sarin, K.Y., Whitson, R.J., Li, J.R., Kim, G., Rezaee, M., Ally, M.S., Kim, J., Yao, C., Chang, A.L., et al. (2015). Smoothened variants explain the majority of drug resistance in basal cell carcinoma. Cancer Cell 27, 342–353. 10.1016/j.ccell.2015.02.002.

11. Chen, M., Gavrilova, O., Zhao, W.Q., Nguyen, A., Lorenzo, J., Shen, L., Nackers, L., Pack, S., Jou, W., and Weinstein, L.S. (2005). Increased glucose tolerance and reduced adiposity in the absence of fasting hypoglycemia in mice with liver-specific Gs alpha deficiency. J Clin Invest 115, 3217–3227. 10.1172/JCI24196.

12. Belteki, G., Haigh, J., Kabacs, N., Haigh, K., Sison, K., Costantini, F., Whitsett, J., Quaggin, S.E., and Nagy, A. (2005). Conditional and inducible transgene expression in mice through the combinatorial use of Cre-mediated recombination and tetracycline induction. Nucleic Acids Res 33, e51. 10.1093/nar/gni051.

13. Youssef, K.K., Lapouge, G., Bouvree, K., Rorive, S., Brohee, S., Appelstein, O., Larsimont, J.C., Sukumaran, V., Van de Sande, B., Pucci, D., et al. (2012). Adult interfollicular tumour-initiating cells are reprogrammed into an embryonic hair follicle progenitor-like fate during basal cell carcinoma initiation. Nature cell biology 14, 1282–1294. 10.1038/ncb2628.

14. Depianto, D., Kerns, M.L., Dlugosz, A.A., and Coulombe, P.A. (2010). Keratin 17 promotes epithelial proliferation and tumor growth by polarizing the immune response in skin. Nature genetics 42, 910–914. 10.1038/ng.665.

15. Joost, S., Zeisel, A., Jacob, T., Sun, X., La Manno, G., Lonnerberg, P., Linnarsson, S., and Kasper, M. (2016). Single-Cell Transcriptomics Reveals that Differentiation and Spatial Signatures Shape Epidermal and Hair Follicle Heterogeneity. Cell Syst 3, 221–237 e229. 10.1016/j.cels.2016.08.010.

16. Quigley, D.A., Kandyba, E., Huang, P., Halliwill, K.D., Sjolund, J., Pelorosso, F., Wong, C.E., Hirst, G.L., Wu, D., Delrosario, R., et al. (2016). Gene Expression Architecture of Mouse Dorsal and Tail Skin Reveals Functional Differences in Inflammation and Cancer. Cell reports 16, 1153–1165. 10.1016/j.celrep.2016.06.061.

17. Ghuwalewala, S., Lee, S.A., Jiang, K., Baidya, J., Chovatiya, G., Kaur, P., Shalloway, D., and Tumbar, T. (2022). Binary organization of epidermal basal domains highlights robustness to environmental exposure. EMBO J 41, e110488. 10.15252/embj.2021110488.

18. Nguyen, M.B., Flora, P., Branch, M.C., Weber, M., Zheng, X.Y., Sivan, U., Joost, S., Annusver, K., Zheng, D., Kasper, M., and Ezhkova, E. (2024). Tenascin-C expressing touch dome keratinocytes exhibit characteristics of all epidermal lineages. Sci Adv 10, eadi5791. 10.1126/sciadv.adi5791.

19. Yang, Y., Gomez, N., Infarinato, N., Adam, R.C., Sribour, M., Baek, I., Laurin, M., and Fuchs, E. (2023). The pioneer factor SOX9 competes for epigenetic factors to switch stem cell fates. Nature cell biology 25, 1185–1195. 10.1038/s41556-023-01184-y.

20. Powell, A.E., Wang, Y., Li, Y., Poulin, E.J., Means, A.L., Washington, M.K., Higginbotham, J.N., Juchheim, A., Prasad, N., Levy, S.E., et al. (2012). The pan-ErbB negative regulator Lrig1 is an intestinal stem cell marker that functions as a tumor suppressor. Cell 149, 146–158. 10.1016/j.cell.2012.02.042.

21. Barker, N., van Es, J.H., Kuipers, J., Kujala, P., van den Born, M., Cozijnsen, M., Haegebarth, A., Korving, J., Begthel, H., Peters, P.J., and Clevers, H. (2007). Identification of stem cells in small intestine and colon by marker gene Lgr5. Nature 449, 1003–1007. 10.1038/nature06196.

22. Weigert, R., Sramkova, M., Parente, L., Amornphimoltham, P., and Masedunskas, A. (2010). Intravital microscopy: a novel tool to study cell biology in living animals. Histochem Cell Biol 133, 481–491. 10.1007/s00418-010-0692-z.

23. Tokita, M.J., Nahas, S., Briggs, B., Malicki, D.M., Mesirov, J.P., Reyes, I.A.C., Farnaes, L., Levy, M.L., Kingsmore, S.F., Dimmock, D., et al. (2019). Biallelic loss of GNAS in a patient with pediatric medulloblastoma. Cold Spring Harb Mol Case Stud 5. 10.1101/mcs.a004572.

24. Skowron, P., Farooq, H., Cavalli, F.M.G., Morrissy, A.S., Ly, M., Hendrikse, L.D., Wang, E.Y., Djambazian, H., Zhu, H., Mungall, K.L., et al. (2021). The transcriptional landscape of Shh medulloblastoma. Nat Commun 12, 1749. 10.1038/s41467-021-21883-0.

25. Bonilla, X., Parmentier, L., King, B., Bezrukov, F., Kaya, G., Zoete, V., Seplyarskiy, V.B., Sharpe, H.J., McKee, T., Letourneau, A., et al. (2016). Genomic analysis identifies new drivers and progression pathways in skin basal cell carcinoma. Nature genetics 48, 398–406. 10.1038/ng.3525.

26. Pedro, M.P., Salinas Parra, N., Gutkind, J.S., and Iglesias-Bartolome, R. (2020). Activation of G-Protein Coupled Receptor-Galphai Signaling Increases Keratinocyte Proliferation and Reduces Differentiation, Leading to Epidermal Hyperplasia. The Journal of investigative dermatology 140, 1195–1203 e1193. 10.1016/j.jid.2019.10.012.

27. Mukhopadhyay, S., Wen, X., Ratti, N., Loktev, A., Rangell, L., Scales, S.J., and Jackson, P.K. (2013). The ciliary G-protein-coupled receptor Gpr161 negatively regulates the Sonic hedgehog pathway via cAMP signaling. Cell 152, 210–223. 10.1016/j.cell.2012.12.026.

28. Cullen, M., Elzarrad, M.K., Seaman, S., Zudaire, E., Stevens, J., Yang, M.Y., Li, X., Chaudhary, A., Xu, L., Hilton, M.B., et al. (2011). GPR124, an orphan G protein-coupled receptor, is required for CNS-specific vascularization and establishment of the blood-brain barrier. Proceedings of the National Academy of Sciences of the United States of America 108, 5759–5764. 10.1073/pnas.1017192108.

29. Shimada, I.S., Hwang, S.H., Somatilaka, B.N., Wang, X., Skowron, P., Kim, J., Kim, M., Shelton, J.M., Rajaram, V., Xuan, Z., et al. (2018). Basal Suppression of the Sonic Hedgehog Pathway by the G-Protein-Coupled Receptor Gpr161 Restricts Medulloblastoma Pathogenesis. Cell reports 22, 1169–1184. 10.1016/j.celrep.2018.01.018.

30. Hwang, S.H., White, K.A., Somatilaka, B.N., Shelton, J.M., Richardson, J.A., and Mukhopadhyay, S. (2018). The G protein-coupled receptor Gpr161 regulates forelimb formation, limb patterning and skeletal morphogenesis in a primary cilium-dependent manner. Development 145. 10.1242/dev.154054.

31. Bai, C.B., Auerbach, W., Lee, J.S., Stephen, D., and Joyner, A.L. (2002). Gli2, but not Gli1, is required for initial Shh signaling and ectopic activation of the Shh pathway. Development 129, 4753–4761.

32. He, X., Zhang, L., Chen, Y., Remke, M., Shih, D., Lu, F., Wang, H., Deng, Y., Yu, Y., Xia, Y., et al. (2014). The G protein alpha subunit Galphas is a tumor suppressor in Sonic hedgehog-driven medulloblastoma. Nature medicine 20, 1035–1042. 10.1038/nm.3666.

33. Eckle, T., Krahn, T., Grenz, A., Kohler, D., Mittelbronn, M., Ledent, C., Jacobson, M.A., Osswald, H., Thompson, L.F., Unertl, K., and Eltzschig, H.K. (2007). Cardioprotection by ecto-5’-nucleotidase (CD73) and A2B adenosine receptors. Circulation 115, 1581–1590. 10.1161/CIRCULATIONAHA.106.669697.

34. Happ, J.T., Arveseth, C.D., Bruystens, J., Bertinetti, D., Nelson, I.B., Olivieri, C., Zhang, J., Hedeen, D.S., Zhu, J.F., Capener, J.L., et al. (2022). A PKA inhibitor motif within SMOOTHENED controls Hedgehog signal transduction. Nat Struct Mol Biol 29, 990–999. 10.1038/s41594-022-00838-z.

35. Pusapati, G.V., Kong, J.H., Patel, B.B., Gouti, M., Sagner, A., Sircar, R., Luchetti, G., Ingham, P.W., Briscoe, J., and Rohatgi, R. (2018). G protein-coupled receptors control the sensitivity of cells to the morphogen Sonic Hedgehog. Science signaling 11. 10.1126/scisignal.aao5749.

36. Somatilaka, B.N., Hwang, S.H., Palicharla, V.R., White, K.A., Badgandi, H., Shelton, J.M., and Mukhopadhyay, S. (2020). Ankmy2 Prevents Smoothened-Independent Hyperactivation of the Hedgehog Pathway via Cilia-Regulated Adenylyl Cyclase Signaling. Developmental cell 54, 710–726 e718. 10.1016/j.devcel.2020.06.034.

37. Youssef, K.K., Van Keymeulen, A., Lapouge, G., Beck, B., Michaux, C., Achouri, Y., Sotiropoulou, P.A., and Blanpain, C. (2010). Identification of the cell lineage at the origin of basal cell carcinoma. Nature cell biology 12, 299–305. 10.1038/ncb2031.

38. Choquet, H., Ashrafzadeh, S., Kim, Y., Asgari, M.M., and Jorgenson, E. (2020). Genetic and environmental factors underlying keratinocyte carcinoma risk. JCI Insight 5. 10.1172/jci.insight.134783.

39. Kilgour, J.M., Jia, J.L., and Sarin, K.Y. (2021). Review of the Molecular Genetics of Basal Cell Carcinoma; Inherited Susceptibility, Somatic Mutations, and Targeted Therapeutics. Cancers (Basel) 13. 10.3390/cancers13153870.

40. Chiang, A., Solis, D.C., Rogers, H., Sohn, G.K., Cho, H.G., Saldanha, G., Lapidus, D., Li, S., Sarin, K.Y., and Tang, J.Y. (2021). Prevalence and risk factors for high-frequency basal cell carcinoma in the United States. J Am Acad Dermatol 84, 1493–1495. 10.1016/j.jaad.2020.07.088.

41. Liu, Y., Banka, S., Huang, Y., Hardman-Smart, J., Pye, D., Torrelo, A., Beaman, G.M., Kazanietz, M.G., Baker, M.J., Ferrazzano, C., et al. (2022). Germline intergenic duplications at Xq26.1 underlie Bazex-Dupre-Christol basal cell carcinoma susceptibility syndrome. The British journal of dermatology 187, 948–961. 10.1111/bjd.21842.

42. Eccles, R.L., Czajkowski, M.T., Barth, C., Muller, P.M., McShane, E., Grunwald, S., Beaudette, P., Mecklenburg, N., Volkmer, R., Zuhlke, K., et al. (2016). Bimodal antagonism of PKA signalling by ARHGAP36. Nat Commun 7, 12963. 10.1038/ncomms12963.

43. Wu, V.H., Yung, B.S., Faraji, F., Saddawi-Konefka, R., Wang, Z., Wenzel, A.T., Song, M.J., Pagadala, M.S., Clubb, L.M., Chiou, J., et al. (2023). The GPCR-Galpha(s)-PKA signaling axis promotes T cell dysfunction and cancer immunotherapy failure. Nat Immunol 24, 1318–1330. 10.1038/s41590-023-01529-7.

44. Chalmers, Z.R., Connelly, C.F., Fabrizio, D., Gay, L., Ali, S.M., Ennis, R., Schrock, A., Campbell, B., Shlien, A., Chmielecki, J., et al. (2017). Analysis of 100,000 human cancer genomes reveals the landscape of tumor mutational burden. Genome Med 9, 34. 10.1186/s13073-017-0424-2.

45. Jayaraman, S.S., Rayhan, D.J., Hazany, S., and Kolodney, M.S. (2014). Mutational landscape of basal cell carcinomas by whole-exome sequencing. The Journal of investigative dermatology 134, 213–220. 10.1038/jid.2013.276.

46. Grachtchouk, V., Grachtchouk, M., Lowe, L., Johnson, T., Wei, L., Wang, A., de Sauvage, F., and Dlugosz, A.A. (2003). The magnitude of hedgehog signaling activity defines skin tumor phenotype. EMBO J 22, 2741–2751. 10.1093/emboj/cdg271.

47. Yang, S.H., Andl, T., Grachtchouk, V., Wang, A., Liu, J., Syu, L.J., Ferris, J., Wang, T.S., Glick, A.B., Millar, S.E., and Dlugosz, A.A. (2008). Pathological responses to oncogenic Hedgehog signaling in skin are dependent on canonical Wnt/beta3-catenin signaling. Nature genetics 40, 1130–1135. 10.1038/ng.192.

48. Tjarks, B.J., Gardner, J.M., and Riddle, N.D. (2019). Hamartomas of skin and soft tissue. Semin Diagn Pathol 36, 48–61. 10.1053/j.semdp.2018.12.001.

49. Xie, J., Murone, M., Luoh, S.M., Ryan, A., Gu, Q., Zhang, C., Bonifas, J.M., Lam, C.W., Hynes, M., Goddard, A., et al. (1998). Activating Smoothened mutations in sporadic basal-cell carcinoma. Nature 391, 90–92. 10.1038/34201.

50. Yuan, Y., Park, J., Feng, A., Awasthi, P., Wang, Z., Chen, Q., and Iglesias-Bartolome, R. (2020). YAP1/TAZ-TEAD transcriptional networks maintain skin homeostasis by regulating cell proliferation and limiting KLF4 activity. Nat Commun 11, 1472. 10.1038/s41467-020-15301-0.

51. Shwartz, Y., Gonzalez-Celeiro, M., Chen, C.L., Pasolli, H.A., Sheu, S.H., Fan, S.M., Shamsi, F., Assaad, S., Lin, E.T., Zhang, B., et al. (2020). Cell Types Promoting Goosebumps Form a Niche to Regulate Hair Follicle Stem Cells. Cell 182, 578–593 e519. 10.1016/j.cell.2020.06.031.

52. Marin-Castejon, A., Marco-Bonilla, M., Terencio, M.C., Arasa, J., Carceller, M.C., Ferrandiz, M.L., Noguera, M.A., Andres-Ejarque, R., and Montesinos, M.C. (2024). Adenosine A(2B) receptor agonist improves epidermal barrier integrity in a murine model of epidermal hyperplasia. Biomed Pharmacother 173, 116401. 10.1016/j.biopha.2024.116401.

53. Cristofalo, V.J., Allen, R.G., Pignolo, R.J., Martin, B.G., and Beck, J.C. (1998). Relationship between donor age and the replicative lifespan of human cells in culture: a reevaluation. Proceedings of the National Academy of Sciences of the United States of America 95, 10614–10619. 10.1073/pnas.95.18.10614.

54. Korsunsky, I., Millard, N., Fan, J., Slowikowski, K., Zhang, F., Wei, K., Baglaenko, Y., Brenner, M., Loh, P.R., and Raychaudhuri, S. (2019). Fast, sensitive and accurate integration of single-cell data with Harmony. Nat Methods 16, 1289–1296. 10.1038/s41592-019-0619-0.

55. McLaren, W., Gil, L., Hunt, S.E., Riat, H.S., Ritchie, G.R., Thormann, A., Flicek, P., and Cunningham, F. (2016). The Ensembl Variant Effect Predictor. Genome Biol 17, 122. 10.1186/s13059-016-0974-4.

56. Melis, N., Subramanian, B., Chen, D., and Weigert, R. (2022). Imaging Neutrophil Migration in the Mouse Skin to Investigate Subcellular Membrane Remodeling Under Physiological Conditions. J Vis Exp. 10.3791/63581.

